# Longitudinal single cell transcriptional and epigenetic mapping of effector, memory, and exhausted CD8 T cells reveals shared biological circuits across distinct cell fates

**DOI:** 10.1101/2022.03.27.485974

**Authors:** Josephine R. Giles, Shin Foong Ngiow, Sasikanth Manne, Amy E. Baxter, Omar Khan, Ping Wang, Ryan Staupe, Mohamed S. Abdel-Hakeem, Hua Huang, Divij Mathew, Mark M. Painter, Jennifer E. Wu, Yinghui Jane Huang, Rishi Goel, E. John Wherry

## Abstract

Naïve CD8 T cells can differentiate into effector (T_EFF_), memory (T_MEM_), or exhausted (T_EX_) CD8 T cells. These developmental pathways are associated with distinct transcriptional and epigenetic changes that endow cells with different functional capacities and therefore therapeutic potential. The molecular circuitry underlying these developmental trajectories and the extent of heterogeneity within T_EFF_, T_MEM_, T_EX_ populations remain poorly understood. Here, we used the lymphocytic choriomeningitis virus model of acutely-resolved and chronic infection and addressed these gaps by applying longitudinal scRNA-seq and scATAC-seq analysis. These analyses uncovered new subsets, including a subpopulation of T_EX_ expressing NK cell-associated genes, as well as multiple distinct TCF1+ stem/progenitor-like subsets in acute and chronic infection. These data also revealed insights into the reshaping of T_EX_ subsets following PD1 pathway blockade and identified a key role for the cell stress regulator, Btg1, in T_EX_ differentiation. Finally, these results highlighted how the same biological circuits such as cytotoxicity or stem/progenitor pathways can be used by CD8 T cells with highly divergent underlying chromatin landscapes. Thus, this transcriptional and chromatin accessibility landscape map elucidates developmental biology and underlying mechanisms governing T_EFF_, T_MEM_, and T_EX_ differentiation.

## Introduction

Upon activation, CD8 T cells can differentiate into effector (T_EFF_) and memory (T_MEM_) T cells following acutely-resolved infections or vaccination, or exhausted T cells (T_EX_) in chronic infections, cancer, and autoimmunity. The importance of transcriptional and epigenetic changes that accompany these developmental choices has been highlighted in recent years (1). However, major questions remain regarding heterogeneity within these populations and how these fate decisions are regulated.

During an acutely-resolving infection or following vaccination, activated CD8 T cells differentiate into T_EFF_ populations that are associated with control of infection and subsequent formation of T_MEM_ that confer long-term protection (2, 3). Among these major branches of CD8 T cell differentiation, many subsets have been identified based on surface phenotype, function, and differentiation potential. For example, combinations of KLRG1, CD127, CX3CR1, and other molecules can identify subsets of T_EFF_ cells with robust effector activity, but limited durability, or alternatively, enhanced capacity to populate the long-term T_MEM_ pool (4, 5). However, how the development of this subset diversity is linked to the underlying transcriptional and epigenetic wiring of these cells remains incompletely understood.

During chronic infection, cancer, and autoimmunity, persistent stimulation induces differentiation of T_EX_. Similar to T_EFF_ and T_MEM_, the T_EX_ population is heterogenous (4, 6). There has been considerable interest in understanding the ontogeny and function of T_EX_ subsets because some subsets are necessary for response to immunotherapies, including to anti-PD1 (7–9) and adoptive T cell therapy (10). Although various definitions have been used, most studies point to the existence of: (i) progenitor T_EX_ (aka “stemlike” or “precursor”); (ii) intermediate or transitory T_EX_; and (iii) terminal T_EX_ (9, 11–14). As a population, T_EX_ cells have a distinct epigenetic landscape compared to T_EFF_ and T_MEM_ (15–18) governed at least in part by the transcription factor (TF) TOX (19–22). Despite many differences, including this divergent epigenetic landscape, T_EX_ share some features with T_EFF_ and T_MEM_. For example, both T_EFF_ and T_EX_ can be cytolytic, although T_EX_ less so, and subsets of T_MEM_ and T_EX_ can have long-term persistence despite different signals for homeostasis (6).

There are key gaps in our understanding of the developmental relationships and transcriptional and epigenetic mechanisms governing T_EFF_, T_MEM_, and T_EX_ differentiation and heterogeneity. These knowledge gaps are due in part to a paucity of paired transcriptional and chromatin accessibility data from CD8 T cells differentiating down these distinct trajectories. It is unclear whether subsets of T_EFF_, T_MEM_, and T_EX_ largely defined using a small number of proteins by flow cytometry reflect underlying cell type heterogeneity. For example, this phenotypic heterogeneity could represent different activation states of the same underlying cell “fate” as defined by chromatin accessibility patterns. Furthermore, some subsets of T_EFF_ and T_MEM_ versus T_EX_ have overlapping protein expression patterns, such as the “progenitor-associated” TF, TCF1. Whether TCF1-expressing cells have the same underlying developmental program or whether TCF1 circuits are used by CD8 T cells arising from different developmental lineages is unclear.

To address these questions, we used the lymphocytic choriomeningitis virus (LCMV) model of acutely-resolved or chronic viral infection to generate longitudinal single cell RNA sequencing (scRNA-seq) and single cell assay for transposase-accessible chromatin sequencing (scATAC-seq) of T_EFF_, T_MEM_, and T_EX_. These data defined population heterogeneity and identified gene expression and accessible chromatin patterns associated with major branches of CD8 T cell differentiation. Comparing scATAC-seq and scRNA-seq revealed that cells with the same accessible chromatin profile existed in more than one transcriptional state. These analyses also uncovered new subpopulations of T_EFF_, T_MEM_, and T_EX_, including a subset of T_EX_ expressing NK cell-associated genes with unique features that may be relevant for immunotherapy. In addition, we found multiple epigenetically distinct populations of TCF1+ antigen-experienced CD8 T cells. T_EX_ precursor cells found early in chronic infection were distinct from T_EX_ progenitors at later timepoints, and both of these TCF1+ populations were different from T_MEM_ precursors and mature T_MEM_ generated after acute infection. Finally, we identify the cell stress response gene, B cell translocation gene (BTG)/TOB family member (BTG1) as a novel regulator of T_EX_ differentiation, highlighting the ability of epigenomic information to uncover new biology and potential therapeutic targets of CD8 T cell differentiation. Thus, this transcriptional and chromatin accessibility landscape map provides insights into the developmental biology and underlying mechanisms governing T_EFF_, T_MEM_, and T_EX_ differentiation.

## Results

### Transcriptional and epigenetic atlas of CD8 T cell differentiation states

To generate a transcriptional and epigenetic longitudinal map of T cell differentiation, we examined virus-specific T cells in acutely-resolved LCMV Armstrong (Arm) infection that generates T_EFF_ and T_MEM_ or LCMV clone 13 (Cl13) infection that results in chronic infection and differentiation of T_EX_ (**Fig. 1a**). We adoptively transferred TCR transgenic gp33-specific (P14) CD8 T cells into congenically distinct recipient mice, infected with Arm or Cl13, then isolated P14 cells from these mice and performed scRNA-seq and scATAC-seq on days 8, 15, and 30 post infection (p.i.) (**Fig. 1a,b**). To visualize the transcriptional and epigenetic T cell landscape, we projected all cells from scRNA-seq or scATAC-seq into UMAP space. This analysis revealed clear separation of cells based on infection (Arm or Cl13) and timepoint (**Fig. 1c, Fig. S1a,b**). scATAC-seq separated cells more clearly, reflecting the enhanced ability of ATAC-seq to distinguish distinct cell types compared to scRNA-seq (23–25). From non-naïve CD8 T cells, we resolved 18 distinct scRNA-seq clusters (**Fig. 1d,e**), and 16 distinct scATAC-seq clusters (**Fig. 1d,f**). Most clusters contained a majority of cells from one infection at one timepoint. However, some clusters were more diverse; scRNA-seq clusters 12-18 contained a mixture of cells from d15 and d30 of Cl13 infection whereas these time points remain more homogenous by scATAC-seq (**Fig. 1e,f**). This latter observation indicates the transcriptional program of T_EX_ is established by d15, but the underlying chromatin landscape of T_EX_ continues to evolve for at least one month.

**Figure 1.**
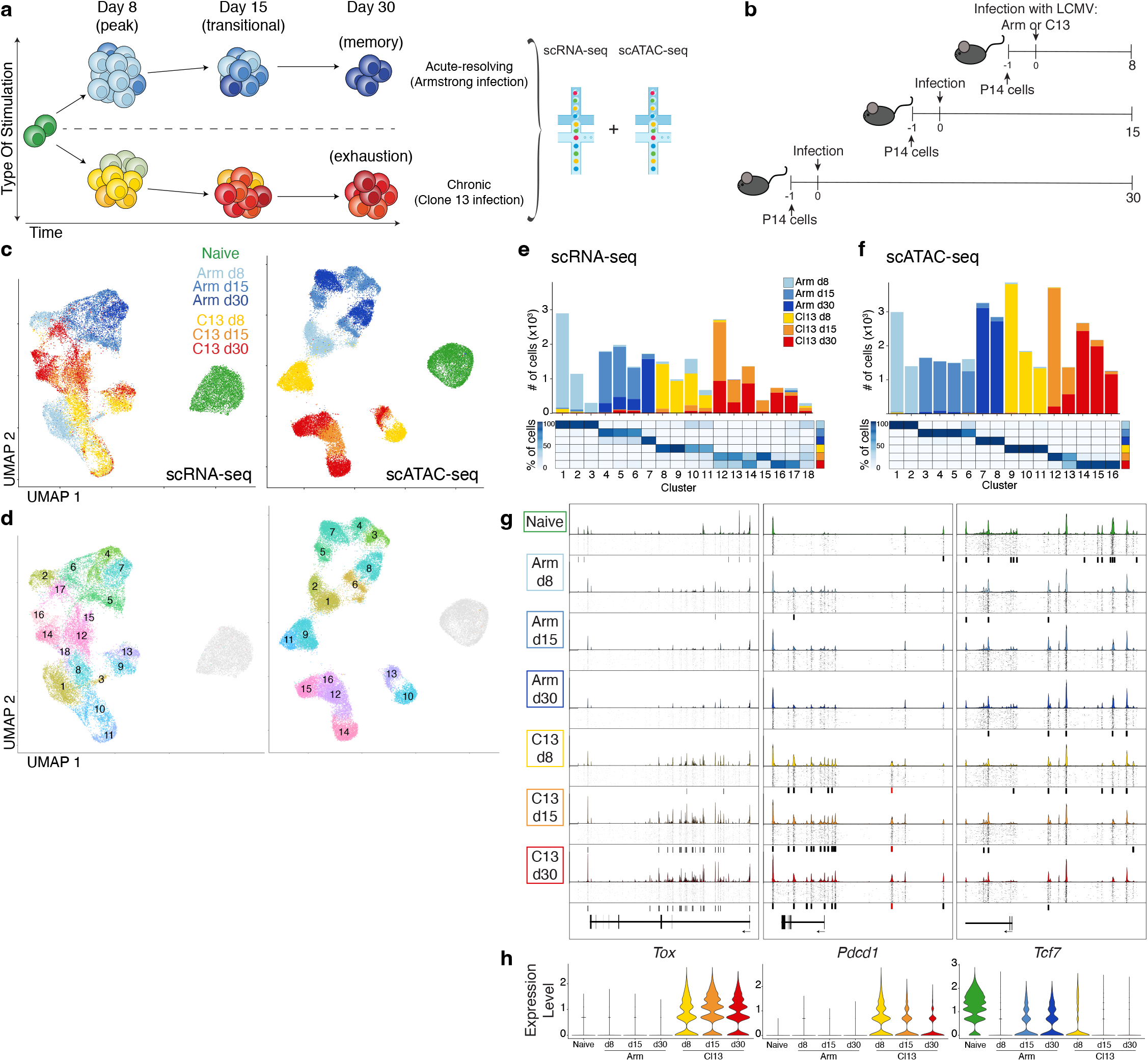
Single cell transcriptional and accessible chromatin landscape of memory and exhausted CD8 T development. **a**) Experimental strategy to capture CD8 T cell differentiation in acute-resolving and chronic viral infection. **b**) Detailed experimental schematic. (**c-d**)UMAP from scRNA-seq (left) and scATAC-seq (right) colored by infection and timepoint (**c**) or by cluster (**d**). (**e-f**) Enumeration and proportion of cells per cluster as indicated for scRNA-seq (**e**) or scATAC-seq (**f**). **g**) scATAC-seq coverage and tile plots. Sample-specific ACRs indicated with black boxes below tile plot. Previously identified *Pdcd1* enhancer (18) indicated in red. **h**) Gene expression from scRNA-seq of genes represented in (**g**).

We next asked how key chromatin accessibility changes identified by scATAC-seq associated with developmental trajectories in Arm or Cl13 infection. We examined 3 canonical CD8 T cell differentiation genes: *Tox, Pdcd1* (encoding PD1), and *Tcf7* (encoding TCF1) (**Fig. 1g**). *Tox*, a TF required for formation of T_EX_, was highly expressed during Cl13 infection at all timepoints (19–22) (**Fig. 1g,h**). The scATAC-seq profiles at the *Tox* locus revealed higher accessibility during Cl13 infection; this accessibility increased over time. *Pdcd1*, encoding the inhibitory receptor PD1, was expressed in Cl13 and had uniquely accessible regions, including a previously described enhancer ~23.8kb upstream of the TSS (15, 18). On the other hand, *Tcf7* was robustly expressed in naïve and lowly expressed at other time points in both infections, reflecting the known expression of this TF following acutely-resolved and chronic infection. Overall accessibility at this locus was greatest in naïve T cells; however, infection- and time-specific accessible chromatin regions (ACRs) suggested complex and varied gene regulation in different populations. Thus, we could identify distinct chromatin accessibility patterns associated with expression of key genes in T_EFF_, T_MEM_, and T_EX_.

### CD8 T cell differentiation following acute infection results in two major lineages delineated by cytotoxic potential

To investigate the developmental trajectories of T_EFF_ and T_MEM_ in acute-resolving infection, we first identified T cell subsets using scRNA-seq (**Fig. 2a**). On d8, three clusters were identified: a memory precursor (MP) cluster, an effector (Eff) cluster, and a cytolytic (CTL) cluster characterized by high expression of *Klrg1, Cx3cr1*, and cytotoxic genes (**Fig. 2b,c**), with the latter likely a subpopulation of the KLRG1+CD127-short-lived effector population (3). At d15, three additional clusters were identified that we named transitional (Trans): Trans I, Trans II, Trans CTL. By d30, there was one primary cluster of memory CD8 T cells (Mem) (**Fig. 2b,c**). We next asked the same question using chromatin accessibility data. We performed unbiased clustering (**Fig. 2d**) then used gene activity, a metric of local gene accessibility, to approximate gene expression and assign differentiation state (**Fig. 2e**). Some clusters defined by scATAC-seq clearly overlapped with transcriptionally-defined clusters, such as d8 Eff and CTL (**Fig. 2e** and **S2a,b**). However, other clusters were only revealed by scATAC-seq, including Mem-CTL, suggesting that chromatin accessibility may provide additional information about patterns of differentiation.

**Figure 2.**
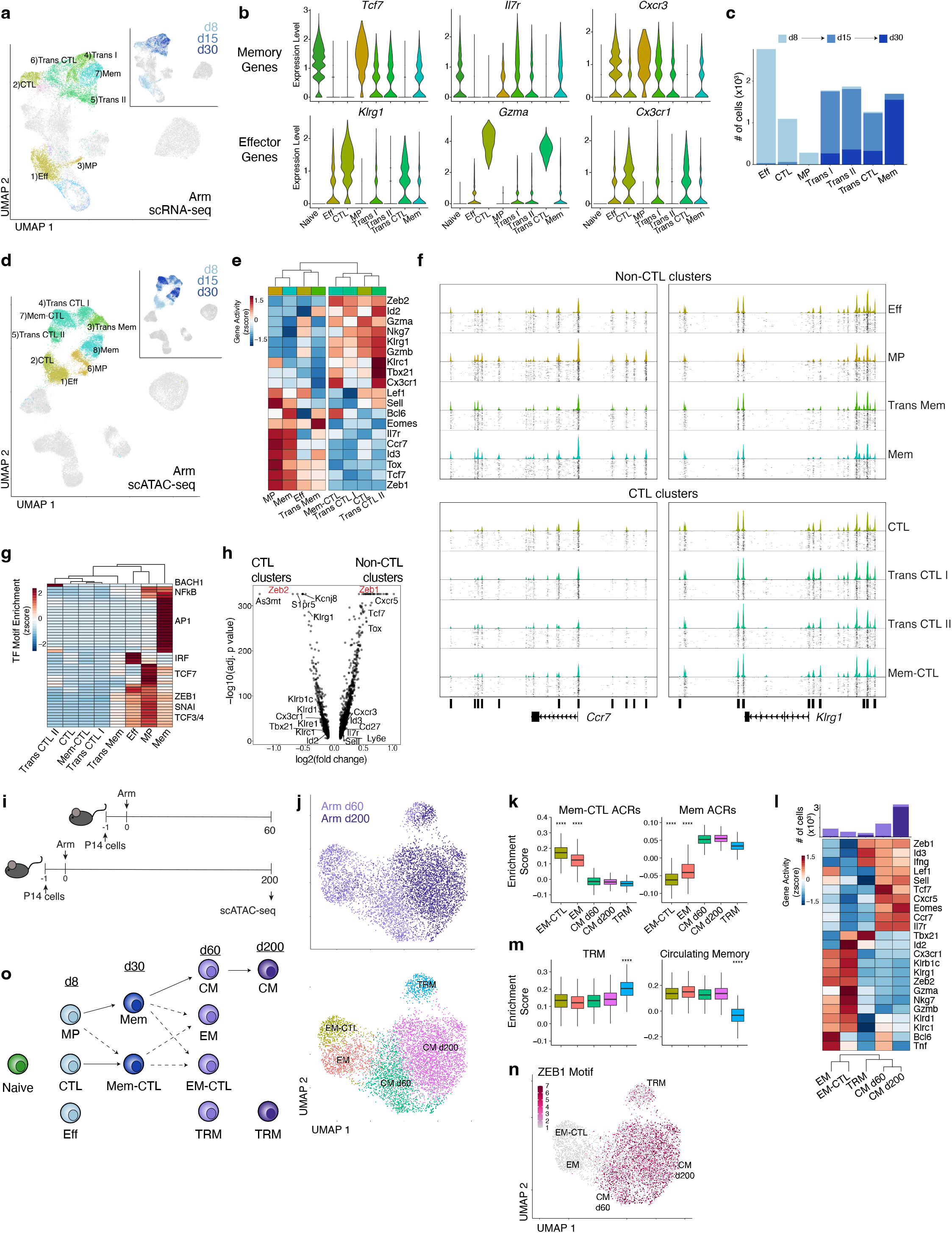
Acute-resolving infection generates two branches of effector and memory CD8 T cells distinguished by epigenetic cytolytic potential. **a**) scRNA-seq UMAP; cells from Arm infection are colored by cluster or timepoint (inset). **b**) Expression of T cell genes by cluster. **c**) Number of cells from Arm infection per cluster filled by timepoint. **d**) scATAC-seq UMAP; cells from Arm infection are colored by cluster or timepoint (inset). **e**) Average gene activity per scATAC-seq cluster; proportion of cells per timepoint in each cluster represented on bottom. **f**) scATAC-seq coverage and tile plots. DACRs of CTL versus non-CTL clusters indicated on the bottom. **g**) Average TF motif enrichment per scATAC-seq cluster of differentially enriched motifs comparing CTL and non-CTL scATAC-seq clusters. **h**) Differential gene activity comparing CTL and non-CTL scATAC-seq clusters. Gene loci of interest indicated. **i**) Experimental schematic of long-term Arm infection experiment. **j**) scATAC-seq UMAP of cells from experiment in (**i**) colored by time point (top) or cluster (bottom). **k**) Enrichment score of cluster-specific ACRs from d30 Arm Mem-CTL and Mem scATAC-seq clusters. Two-sided Wilcoxon test of EM or EM-CTL cells versus all, **** p<0.0001. **l**) Average gene activity per scATAC-seq cluster with number of cells per cluster indicated on top, filled by timepoint. **m**) Enrichment score of gene activity from gene sets derived from TRM or circulating memory cells (57). Wilcoxon test of TRM cells versus all, **** p<0.0001. **n**) scATAC-seq UMAP of cells from experiment in (**i**) colored by ZEB1 motif enrichment. **o**) Data summary schematic. All boxplots represent: center line, median; box limits, upper and lower quartiles; whiskers, 1.5x interquartile range.

Gene activity analysis also revealed two broad epigenetic groups among the eight scATAC-seq clusters that differed in accessibility at cytotoxic genes including *Gzma, Gzmb*, and *Klrc1* (NKG2A): CTL and non-CTL (**Fig. 2e**). CTL clusters included CTL from d8, Trans CTL I and Trans CTL II from d15, and the Mem-CTL cluster from d30. The non-CTL clusters included Eff and MP from d8, Trans Mem from d15, and Mem from d30. These two groups of clusters displayed different ACR profiles including ACRs at the *Ccr7* locus for the non-CTL clusters and *Klrg1* locus for the CTL clusters (**Fig. 2f**). Notably, although there were two distinct clusters of d30 memory CD8 T cells based on chromatin accessibility (Mem and Mem-CTL), there was only one major transcriptional cluster (**Fig. 2a,d, S2a-c**). In summary, scATAC-seq identified two epigenetically distinct groups following acutely-resolved infection that were defined by cytotoxic or memory patterns. This bifurcation was identifiable by d8, consistent with the notion of early commitment to either the memory or effector lineage (2, 26).

To interrogate what TF(s) might regulate these two epigenetic groups, we identified TF binding motifs with differential enrichment in CTL versus non-CTL clusters. Some TFs were more specific for one cluster such as AP1 motifs in the Mem cluster, but ZEB1, TCF3 (E2A), TCF12 (HEB), and SNAI motifs were all enriched in non-CTL clusters (Eff, MP, Trans Mem, Mem) compared to CTL clusters (CTL, Trans CTL I, Trans CTL II, Mem-CTL) (**Fig. 2g**). Based on gene activity, *Zeb1* was likely to be highly expressed in the non-CTL cluster group and *Zeb2* in the CTL clusters (**Fig. 2h**). Although ZEB2 does not have a unique motif that can be tested for enrichment, the ZEB1 motif was strongly enriched in non-CTL clusters and nearly absent in CTL clusters (**Fig. S3**). This analysis is consistent with previous work identifying a role for *Zeb1* in T_MEM_ formation and function whereas *Zeb2* promoted the formation of short-lived T_EFF_ (27–29). However, our data suggest that *Zeb2* may have a specific role in subsets with cytotoxic function, including Mem-CTL cells at d30 and highlight the *Zeb1-Zeb2* TF pair in the bifurcation of CTL and non-CTL branches of CD8 T cell differentiation in acutely-resolved infection.

It was possible that Mem-CTL (and Trans CTL I, Trans CTL II) represented residual CTL effectors that persisted after viral clearance. To test this idea, we next examined whether both Mem and Mem-CTL were present at later time points. We performed scATAC-seq of virus-specific CD8 T cells isolated on days 60 (d60) and 200 (d200) after Arm infection (**Fig. 2i**). Indeed, at d60, two clusters had enriched accessibility at loci associated with the d30 Mem-CTL cluster, which we named effector memory (EM) and EM-CTL (**Fig. 2j-k**). One of these two clusters, EM-CTL, had increased accessibility at cytotoxic gene loci, including *Gzma, Gzmb*, and NK receptors (**Fig. 2l**). Tissue-resident memory (TRM) and central memory (CM) clusters were also present at d60 (**Fig. 2j-m**). However, by d200 most cells belonged to a single CM cluster (CM d200) with a small proportion of TRM cells (**Fig. j-m**). CM cells from d60 and d200 separated into different clusters suggesting continued evolution of the memory CD8 T cell chromatin accessibility landscape over time (**Fig. S4**). TF motif enrichment revealed a clear increase in ZEB1 motif accessibility in CM and TRM cells and a relative absence in EM and EM-CTL (**Fig. 2n**). These data confirm that CD8 T cells similar to the d30 Mem-CTL cluster are also present a month later but are essentially undetectable by d200, consistent with the evolution of the CD8 T cell memory pool to largely CM over time (26) together with TRM (30). Altogether, these data define a trajectory of CD8 T cell differentiation from naïve cells to long-term memory that develops after acute infection (**Fig. 2o**) and suggest that effector functions in longer-lived CD8 T cells may be epigenetically encoded early during infection.

### Temporal scRNA-seq reveals transcriptional heterogeneity of T_EX_

Unlike acute-resolving infections, chronic infection and cancer induce differentiation of T_EX_ which are characterized by expression of IRs and altered effector functions (6). Multiple subsets have been identified within the T_EX_ population, including exhausted progenitor cells, exhausted intermediate cells, and terminally exhausted cells (9, 11–14). The overall heterogeneity in T_EX_, however, remains incompletely understood. To examine the development and heterogeneity of T_EX_ over time, we first defined CD8 T cell clusters from Cl13 infection with scRNA-seq data (**Fig. 3a**) using differential gene expression (**Fig. 3b**), and cluster similarity analysis (**Fig. 3c**). At d8, there were 4 major clusters (**Fig. 3a-c**). One cluster contained effector-like (Eff-like) cells and was distinct from the Eff cluster generated from Arm infection. Among differentially expressed genes (DEGs) between the Cl13 Eff-like and Arm Eff cells, *Tox, Lag3, Rgs16*, and *Ifi27I2a*, were higher in Cl13 Eff-like cells whereas *Klrg1, Ccr2*, and *Selplg* were increased in Eff cells from Arm (**Fig. 3d**). Pathway analysis revealed increased expression of genes involved in general T cell activation in Arm Eff cells, whereas Eff-like cells from Cl13 had increased expression of viral response genes, including those involved in type I interferon (IFN) production (**Fig. 3e**). There were also two proliferating clusters (**Fig. 3b,c,f**). Because cell cycle genes can obscure the underlying transcriptional identity of proliferating cells, we projected these cells back onto the remaining clusters (See Methods). Most proliferating cells belonged to the d8 Eff-like cluster as expected, though a smaller number were derived from clusters present at later time points (**Fig. 2g**) consistent with ongoing cell cycle in T_EX_ populations (14). The fourth d8 Cl13 cluster, Exh-Pre, had some similarity to MP from Arm infection including expression of *Il7r, Id3, Tcf7, Lef1, Sell*, and *Ccr7* (**Fig. 3b,c**). However, this subset also expressed exhaustion-related genes (*Tox*, *Tox2, Pdcd1, Lag3*), confirming previous work that identified an exhaustion-committed population early during Cl13 infection (22, 31).

**Figure 3.**
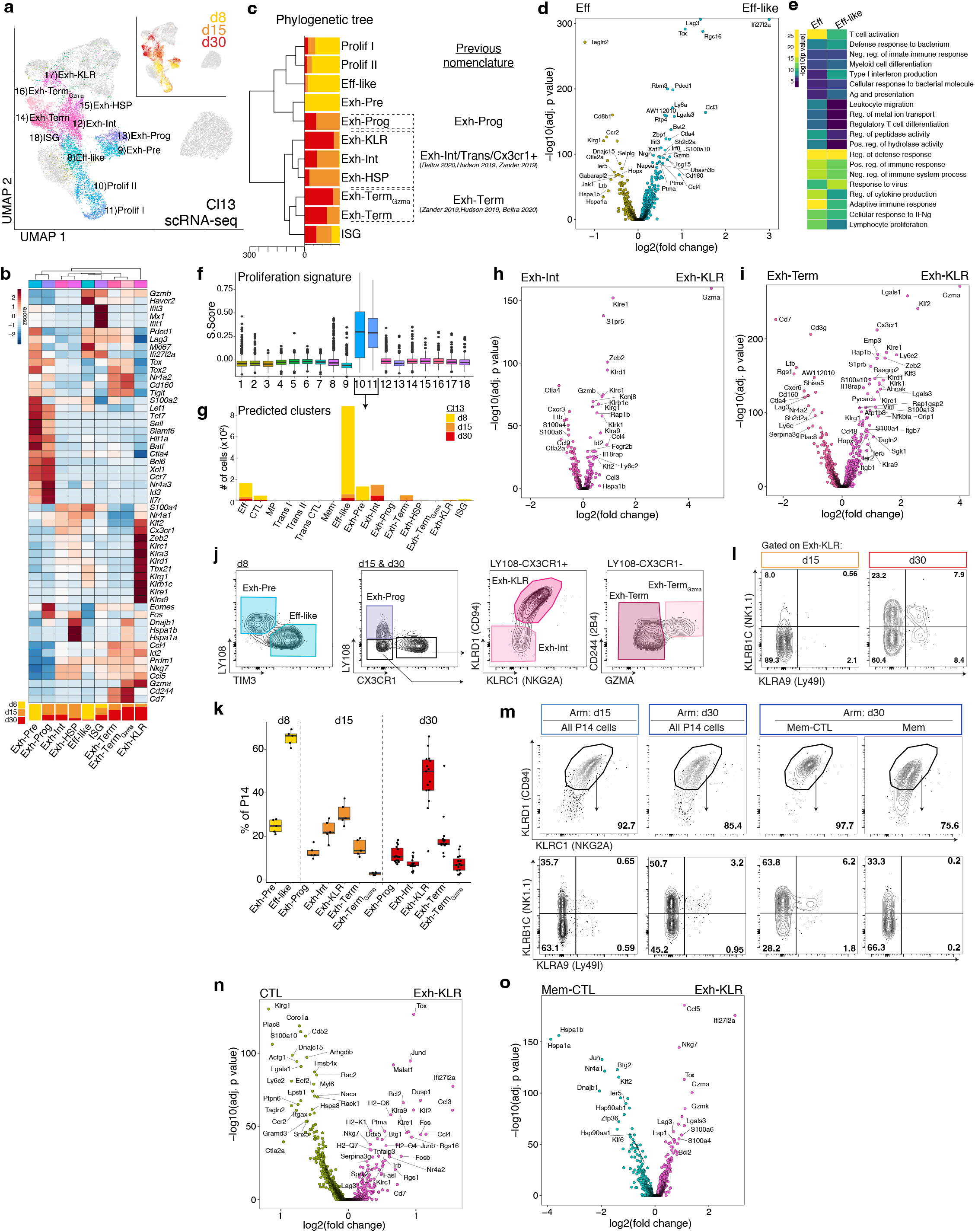
Exhausted CD8 T cells are transcriptionally heterogenous including a distinct subset characterized by expression of NK receptors. **a**) scRNA-seq UMAP; cells from Cl13 infection are colored by cluster or timepoint (inset). **b**) Average gene expression per scRNA-seq cluster with proportion of cells per timepoint in each cluster represented on bottom. **c**) Phylogenetic tree of scRNA-seq clusters with proportion of cells per timepoint. Correspondence of clusters with previous nomenclature indicated. **d**) DEGs between Eff and Eff-like clusters. **e**) Gene ontology analysis of DEGs in (**d**). **f**) Cell cycle S.Score for each cluster (See Methods). Boxplots represent: center line, median; box limits, upper and lower quartiles; whiskers, 1.5x interquartile range, dots, outliers. **g**) Predicted cluster identity of proliferating cells shown as number of cells per cluster and colored by timepoint (See Methods). **h**) DEG analysis between Exh-Int and Exh-KLR clusters. **i**) DEG analysis Exh-Term and Exh-KLR clusters. **j**) Flow cytometry gating strategy to identify T_EX_ clusters. Cells are gated as live single CD8^+^ P14 cells. **k**) Enumeration of T_EX_ clusters gated in (**j**). **l**) Representative flow cytometry plots gated on Exh-KLR cells as in (**j**) from Cl13 infection at d15 and d30, mean percentage per quadrant in bold. **m**) Representative flow cytometry plots from Arm infection at d15 or d30 gated on live singlets CD8^+^ P14 cells (top) or KLRC1^+^KLRD1^+^ P14 cells (bottom) as indicated. Mean percentage per quadrant in bold. All flow cytometry is representative of at least two experiments with 5-10 mice per group. Boxplots represent: center line, median; box limits, upper and lower quartiles; whiskers, 1.5x interquartile range. Each dot represents a mouse. **n**) DEG analysis between CTL cluster from Arm infection and Exh-KLR cluster from Cl13 infection. **o**) DEG analysis between Mem-CTL cluster from Arm infection and Exh-KLR cluster from Cl13 infection. All boxplots represent: center line, median; box limits, upper and lower quartiles; whiskers, 1.5x interquartile range.

We next investigated heterogeneity within the T_EX_ population at later time points when exhaustion is established. Seven clusters were present at d15 and d30 p.i. (**Fig. 3a-c**). An Exh-Prog cluster at these time points was most similar to the d8 Exh-Pre cluster (**Fig. 3c**), but had unique features including high expression of *Eomes* and *Fos* (**Fig. 3b**). The two smallest clusters (**Fig. 1e**) were defined by expression of heat-shock protein genes (Exh-HSP) or interferon-stimulated genes (ISG) (**Fig. 3b**). Two additional clusters with the highest IR expression were likely analogous to previously described terminal T_EX_ subset (9, 11–13); however, unbiased clustering separated these terminal T_EX_ cells into two subsets, Exh-Term and Exh-Term_Gzma_, the latter distinguished by high expression of cytotoxic genes, including *Gzma* (**Fig. 3b,c**). These analyses also revealed a previously unappreciated population of T_EX_ that expressed NK-associated genes, Exh-KLR (**Fig. 3c**). Given the close relationship between Exh-Int, Exh-KLR, and Exh-HSP (**Fig. 3c**), these subsets were likely included in the intermediate exhausted population in previous studies (9, 11–14). To gain more insight into this Exh-KLR subset, we directly compared Exh-KLR cells to Exh-Int (**Fig. 3h**) and Exh-Term (**Fig. 3j**). In both comparisons, Exh-KLR were distinguished by genes associated with NK cells (*Klrs, Fcgr2b*, etc.), cytotoxic genes (*Gzma*, *Gzmb*), migration-related genes (*S1pr5, Itgb7*) and several TFs (*Zeb2, Klf2, Klf3, Id2*). These results suggested that Exh-KLR cells may have more cytolytic potential than other T_EX_ subsets. Recent work has identified potential clinically relevant T cells expressing NK receptors (32–35). T_EX_ with characteristics of this Exh-KLR population have not been previously described, but this subset may have relevance to CD8 T cells with NK-like gene expression in other settings.

Next, we wanted to ask whether these T_EX_ subpopulations, including the Exh-KLR subset, could be identified by flow cytometry. Gating on virus-specific P14 CD8 T cells, at d8, Exh-Pre and Eff-like could be distinguished using LY108 and TIM3 (**Fig. 3j**). At d15 and d30, the major subsets were identified using a tiered gating strategy (**Fig. 3j**). Exh-Prog were LY108+CX3CR1-. From the LY108-CX3CR1+ gate, Exh-KLR were identified by expression of KLRC1 (NKG2A) and KLRD1 (CD94) whereas Exh-Int were NKG2A^-^ CD94^-^. Exh-Term and Exh-Term_GzmA_ from the LY108-CX3CR1-gate were then separated based on GZMA expression. Consistent with the transcriptional data, Exh-Term_Gzma_ cells also had higher expression of CD244 (2B4) (**Fig. 3b,j**). Thus, based on surface phenotype, it was possible to resolve distinct populations of Exh-Pre, Eff-like, Exh-Prog, Exh-Int, Exh-KLR, Exh-Term, and Exh-Term_Gzma_ during chronic viral infection. The relative proportion of these subsets changed over time, with a notable increase in Exh-KLR cells from day 15 to day 30 (**Fig. 3k**). By RNA, the Exh-KLR subset expressed several additional NK receptors, including KLRB1C (NK1.1) and KLRA9 (Ly49I) (**Fig. 3l**). Expression of these proteins increased over time and by day 30 there was heterogeneity within the Exh-KLR population based on combinations of these NK receptors (**Fig. 3l**) perhaps reflecting functional diversification since some of these molecules can recognize stress ligands on infected or transformed cells (36). Expression of NK receptors is not unique to Cl13 and most virus-specific CD8 T cells from Arm-infected mice also expressed KLRC1 and KLRD1 at d8 and d30 and also had variable expression of KLRB1C and KLRA9 (**Fig. 3m**), consistent with early studies documenting expression of NK receptors on CD8 T cells in viral and bacterial infections (37, 38). Given the shared expression of NK receptors and effector-like transcriptional profile, we next directly compared Exh-KLR from Cl13 infection with CTL (**Fig. 3n**) and Mem-CTL (**Fig. 3o**) subsets from d8 and d30 of Arm infection, respectively. Both comparisons revealed a large number of DEGs between Exh-KLR and the two cytolytic cell types from Arm infection, including higher expression of *Tox*, *Bcl2*, and *Lag3* in Exh-KLR. These results indicate that Exh-KLR cells were distinct from T_EFF_ and T_MEM_ CD8 T cells generated from an acute infection (**Fig. 3n,o**). These observations also suggest that despite divergent differentiation of Exh-KLR in chronic infection compared to CTL and Mem-CTL in acute infection, these cells share a transcriptional module containing NK-associated genes. This NK and cytotoxicity module may have functional utility in multiple disease contexts.

### Single cell epigenetic profiling reveals 4 distinct T_EX_ subsets

CD8 T cell exhaustion is the result of an epigenetically distinct developmental path compared to T_EFF_ and T_MEM_ (15, 39), driven at least in part by the TF TOX (19–22). However, it has been unclear how phenotypic or transcriptional heterogeneity within the T_EX_ population is related to underlying chromatin landscape heterogeneity. This is a key question since the distinct epigenetic landscape of T_EX_ limits plasticity and susceptibility to therapeutic manipulation (15–17). To address this question, we next asked whether distinct T_EX_ subsets also existed based on scATAC-seq profiles (**Fig. 4a**).

**Figure 4.**
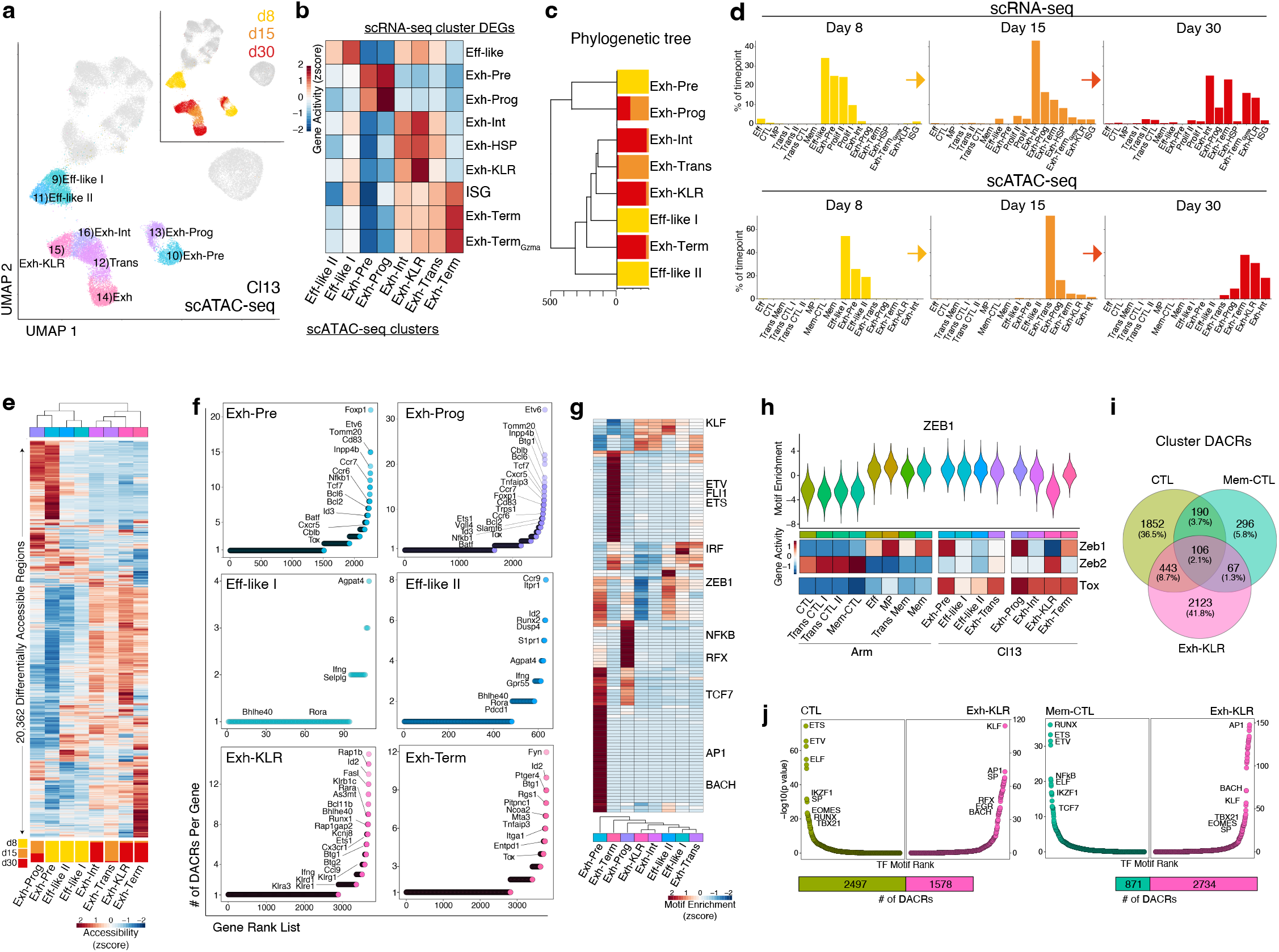
The accessible chromatin landscape of T_EX_ evolves over time and distinguishes fewer epigenetic cell fates under wider transcriptional diversity. **a**) scATAC-seq UMAP; cells from Cl13 infection are colored by cluster or timepoint (inset). **b**) Average enrichment score per scATAC-seq cluster of gene sets from scRNA-seq cluster DEGs, using gene activity. **c**) Phylogenetic tree of scATAC-seq clusters with proportion of cells per timepoint. **d**) Percentage of cells from Cl13 infection by timepoint as indicated in scRNA-seq clusters (top) and scATAC-seq clusters (bottom). **e**) Average accessibility of all DACRs per scATAC-seq cluster with proportion of cells per timepoint in each cluster represented on bottom. **f**) Number of DACRs per gene loci for each scATAC-seq cluster. **g**) Average TF motif enrichment per cluster of differentially enriched TF motifs. **h**) Top, ZEB1 motif enrichment. Bottom, Zeb1, Zeb2, and Tox average gene activity per scATAC-seq cluster. **i**) Venn diagram of overlapping DACRs in scATAC-seq CTL, Mem-CTL, and Exh-KLR clusters. **j**) TF motif enrichment in DACRs comparing scATAC-seq Exh-KLR and CTL (left) or Mem-CTL (right) clusters with total number of DACRs represented on bottom.

Unbiased clustering of scATAC-seq identified 8 clusters of virus-specific CD8 T cells during Cl13 infection. To infer cell subset identity, we used a combination of gene activity (**Fig. 4b**), timepoint, and cluster similarity (**Fig. 4c**). First, we calculated enrichment of gene sets derived from the scRNA-seq clusters (**Fig. 4b**). On d8, there were three clusters: an Exh-Pre cluster and two Eff-like clusters. Direct comparison of Eff-like I and Eff-like II revealed increased accessibility at several genes related to migration in Eff-like II, including *Ccr9, S1pr1, Cd69*, and several integrins (**Fig. S5**), suggesting these cells may traffic to peripheral sites. By d15, the Exh-Prog subset was identifiable by scATACseq, however, the majority of cells were in a second cluster we called transitory (Exh-Trans) because it was almost exclusively found at d15. By d30, most cells populated four clusters that, based on gene activity signatures, we named Exh-Prog, Exh-Int, Exh-KLR, and Exh-Term. To understand how these epigenetically-defined subsets mapped to transcriptionally-defined subsets, we compared scRNA-seq clusters and scATAC-seq clusters by time point (**Fig. 4d**). scATAC-seq resolved fewer clusters than scRNA-seq at d8 (3 clusters vs 7 clusters), d15 (5 vs 10), and d30 (5 vs 11). These differences likely reflect a smaller number of cell “fates” revealed by scATAC-seq underlying multiple transcriptional states.

Next, we investigated epigenetic programs used by different T_EX_ subsets. First, we visualized all 20,362 differentially accessible chromatin regions (DACRs) (**Fig 4e**), then assessed the number of DACRs in each gene locus (**Fig 4f**). This analysis revealed global patterns of shared and distinct ACRs among the T_EX_ clusters. For example, Exh-Pre and Exh-Prog DACR profiles were most similar to each other (**Fig. 4e**), including DACRs at genes associated with stem-like biology such as *Tcf7, Foxp1*, and *Id3* (**Fig. 4f**). The Eff-like I and Eff-like II clusters from chronic infection shared accessibility at *Ifng* and *Bhlhe40*, as did Exh-KLR (**Fig. 4f**). However, Exh-KLR also contained DACRs at *Rap1b, Id2* and *Klrb1c*. Exh-Term exhibited a distinct ACR profile (**Fig. 4e**) that included accessibility at *Fyn, Ptger4, Btg1*, and *Rgs1* (**Fig. 4f**).

We next asked which TFs had the potential to regulate transcriptional programs within each T_EX_ subset (**Fig. 4g**). As expected, ACRs in both Exh-Pre and Exh-Prog were enriched in TCF1 motifs. However, Exh-Pre had a relative increase in accessibility of AP1 motifs, suggesting response to high TCR stimulation, whereas Exh-Prog were more enriched in NFkB and RFX motifs (**Fig. 4 g**). Exh-Term also had a distinct TF motif profile characterized by high enrichment in ETV and ETS TF motifs, including Fli1. Exh-KLR and Exh-Int clusters shared enrichment for several TF binding motifs including the KLF family. However, the Exh-KLR cluster was distinguished from all other clusters by relative absence of ZEB1 motifs (**Fig. 4g,h**), a pattern reminiscent of CTL clusters generated following Arm infection (**Fig. 2g** and **S3**). Furthermore, high *Zeb2* but low *Zeb1* gene activity was also a characteristic of Exh-KLR and CTL clusters from Arm infection, suggesting overlapping TF circuits (**Fig. 4h**). Despite this shared Zeb2-associated “CTL” feature, the Exh-KLR cluster generated during chronic infection had high *Tox* gene activity, which was absent from the CTL clusters from Arm infection (**Fig. 4h**).

To further compare the epigenetic programs used in Exh-KLR and the Arm-derived CTL clusters, we assessed the overlap of cluster-specific ACRs from CTL, Mem-CTL, and Exh-KLR (**Fig. 4i**). Indeed, most Exh-KLR ACRs (2123/2739) were unique compared to Arm-derived CTL and Mem-CTL. In fact, Exh-KLR shared only ~11% and ~3% ACRs with CTL and Mem-CTL, respectively. DACRs unique to Exh-KLR were enriched in KLF and AP1 motifs, whereas those in CTL and Mem-CTL were enriched for ETS, ETV, RUNX motifs (**Figs. 4j**). Thus, the Exh-KLR subset employs epigenetic and transcriptional modules related to cytolytic activity and NK biology that are also used by CD8 T cells in acute-resolving viral infections, but the Exh-KLR subset is otherwise largely distinct from CTL generated during Arm infection.

Collectively, these scATAC-seq data resolved accessible chromatin landscape relationships of pre-T_EX_ and established T_EX_ CD8 T cells, revealed similarities and differences compared to T_EFF_ and T_MEM_ developmental trajectories, and identified 4 major T_EX_ clusters – Exh-Prog, Exh-Int, Exh-KLR, and Exh-Term – based on single cell chromatin accessibility.

### PD1 blockade promotes differentiation of exhausted CD8 T cell subsets

Several recent studies have demonstrated that the distinct epigenetic landscape of T_EX_ limits their ability to re-differentiate following PD1 blockade or removal of antigen (15–17). PD1 blockade targets the Exh-Prog subset (7–9) resulting in an increase in T_EX_ intermediate cells (11, 12). Our data above indicate that this T_EX_ intermediate population is heterogeneous and contains Exh-Int and Exh-KLR subset, but how PD1 blockade impacts the balance of these subsets is unknown. Thus, to investigate how PD1 blockade impacts T_EX_ population dynamics and differentiation, we treated Cl13 infected mice with αPDL1 for two weeks and examined responding T_EX_ by scATAC-seq (**Fig. 5a,b**).

**Figure 5.**
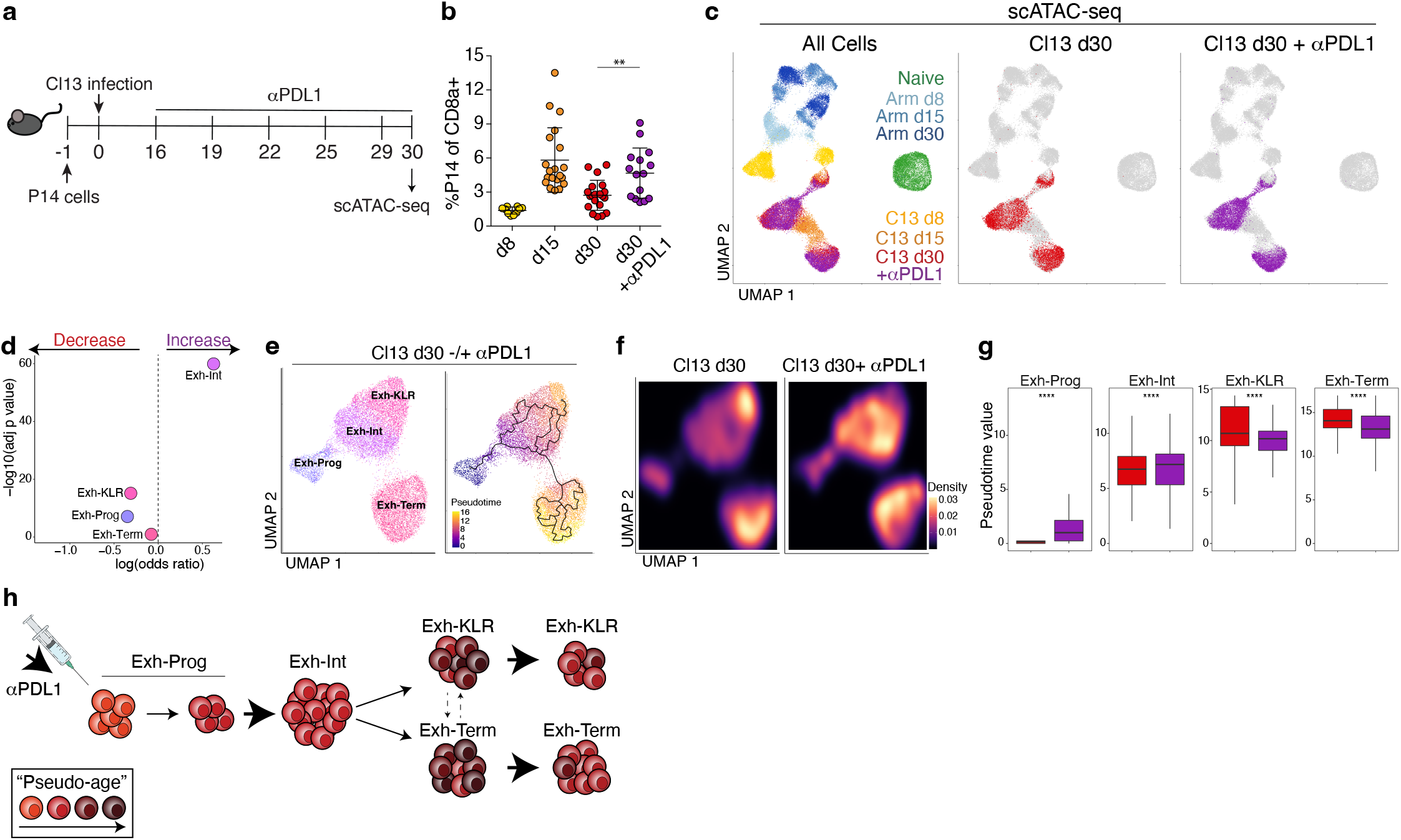
PD1 pathway blockade alters T_EX_ subset dynamics within the pre-existing population structure. **a**) Experimental schematic. **b**) Frequency of blood P14 cells determined by flow cytometry. Day 30 with and without αPDL1 were compared with a two-sided t-test, ** p<0.01 **c**) scATAC-seq UMAPs colored by infection, timepoint, and treatment as indicated. **d**) Difference in the number of cells in each scATAC-seq T_EX_ cluster with and without αPDL1 using Fischer’s exact test. **e**) scATAC-seq UMAP of Cl13 d30 cells with or without αPDL1, colored by cluster (left) or pseudotime calculated by Monocle (right). **f**) scATAC-seq UMAP of Cl13 d30 cells with and without αPDL1 colored by density. **g**) Pseudotime within each scATAC-seq cluster comparing cells with or without αPDL1 using two-sided Wilcoxon test, **** p<0.0001. **h**) Schematic summary of results. All boxplots represent: center line, median; box limits, upper and lower quartiles; whiskers, 1.5x interquartile range.

We first determined where cells from αPDL1-treated mice were positioned in the overall scATAC-seq UMAP space of virus-specific CD8 T cells from Arm and Cl13 infections (**Fig. 5c**). This analysis demonstrated αPDL1-treated cells largely overlapped with T_EX_ from control-treated mice (**Fig. 5c**). PD1 blockade did not produce cells that overlapped with T_EFF_ or T_MEM_ from Arm infection, nor did it result in the formation of a new epigenetic cluster of T_EX_. We next assessed whether αPDL1 altered the frequencies of T_EX_ subsets (**Fig. 5d**). Indeed, PD1 blockade resulted in a substantial increase in Exh-Int cells and a small decrease in Exh-KLR and Exh-Prog clusters; however, these changes were associated with minimal DACRs changes within each subset. This result suggested that αPDL1 affected the trajectory of T_EX_ differentiation by altering the subset distribution augmenting Exh-Int at the expense of Exh-Prog and Exh-KLR. To further investigate this question, we used pseudotime analysis of scATAC-seq data using only cells from d30 of Cl13 infection with and without αPDL1 treatment (**Fig. 5e**). These data illustrate a trajectory from Exh-Prog to Exh-Int and then to either Exh-Term or Exh-KLR (**Fig. 5f**). These data also revealed a clear shift in cell density in UMAP space within these clusters following PD1 pathway blockade (**Fig. 5f**). These analyses revealed an increase in the “pseudo-age” of Exh-Prog and Exh-Int after αPDL1 and a decrease in the “pseudo-age” of Exh-KLR and Exh-Term cells suggesting new cells entered these clusters and/or “older” terminally differentiated T_EX_ were lost after treatment (**Fig. 5g**). Thus, the major effect of PD1 pathway blockade is to accelerate differentiation of Exh-Prog to Exh-Int (**Fig. 5h**). These data also identify a specific rebalancing after αPDL1 with relative depletion of Exh-KLR cells in favor of the Exh-Int subset. Together, these data demonstrate that PD1 pathway blockade alters T_EX_ subset dynamics within the preexisting T_EX_ population hierarchy, an observation with implications for functional and therapeutic impact of PD1 pathway blockade.

### TCF1+ precursors initiate two distinct developmental pathways towards induction of T_MEM_ or T_EX_

A major unresolved question is how similar T_EX_ are compared to CD8 T cells generated following acutely-resolved infection. One such question centers on cells expressing TCF1 (*Tcf7*) that are thought to have progenitor or stem-like activity (40). We therefore compared the *Tcf7*-expressing subsets generated in Arm and Cl13 infection (**Fig. S6**). First, we constructed a phylogenetic tree using all scRNA-seq clusters (**Fig. 6a**). This analysis revealed transcriptional similarity of some subsets at d8 between the two infections; however, by d15, subsets from the same infection were most similar to each other. For example, *Tcf7*-expressing MP from d8 Arm and Exh-Pre from d8 Cl13 infection were transcriptionally similar to each other; however, by d15, subsets from Arm infection formed a unique branch whereas the Exh-Prog subset from Cl13 branched off from d8 Exh-Pre and MP (**Fig. 6a**). UMAP analysis also reflected these relationships where MP, Exh-Pre, and Exh-Prog subsets clustered together in a different UMAP location than either Mem and Naïve cells (**Fig. 6b**). Analysis of DEGs revealed both shared and distinct patterns of gene expression and highlighted the relative quiescence of Mem cells (**Fig. 6c**). Whereas scRNA-seq clustered Exh-Pre, Exh-Prog, and MP together, the scATAC-seq phylogenetic tree revealed Exh-Pre and Exh-Prog were epigenetically distinct from all other clusters, forming their own independent branch (**Fig. 6d**). Also, in contrast to the scRNA-seq data, MP and Mem were most similar to each other based on scATAC-seq (**Fig 6d**). These epigenetic relationships between *Tcf7+* subsets were also clear in a UMAP constructed from scATAC-seq data; Exh-Pre and Exh-Prog colocalized, distal to where MP and Mem colocalized (**Fig. 6e**). These four *Tcf7+* CD8 T cell subsets also displayed distinct chromatin accessibility profiles that highlighted an exhaustion-versus memory-associated ACR pattern (**Fig. 6f**). Together, these data demonstrate epigenetic divergence between virus-specific CD8 T cells in settings that result in T_EX_ versus T_MEM_ differentiation.

**Figure 6.**
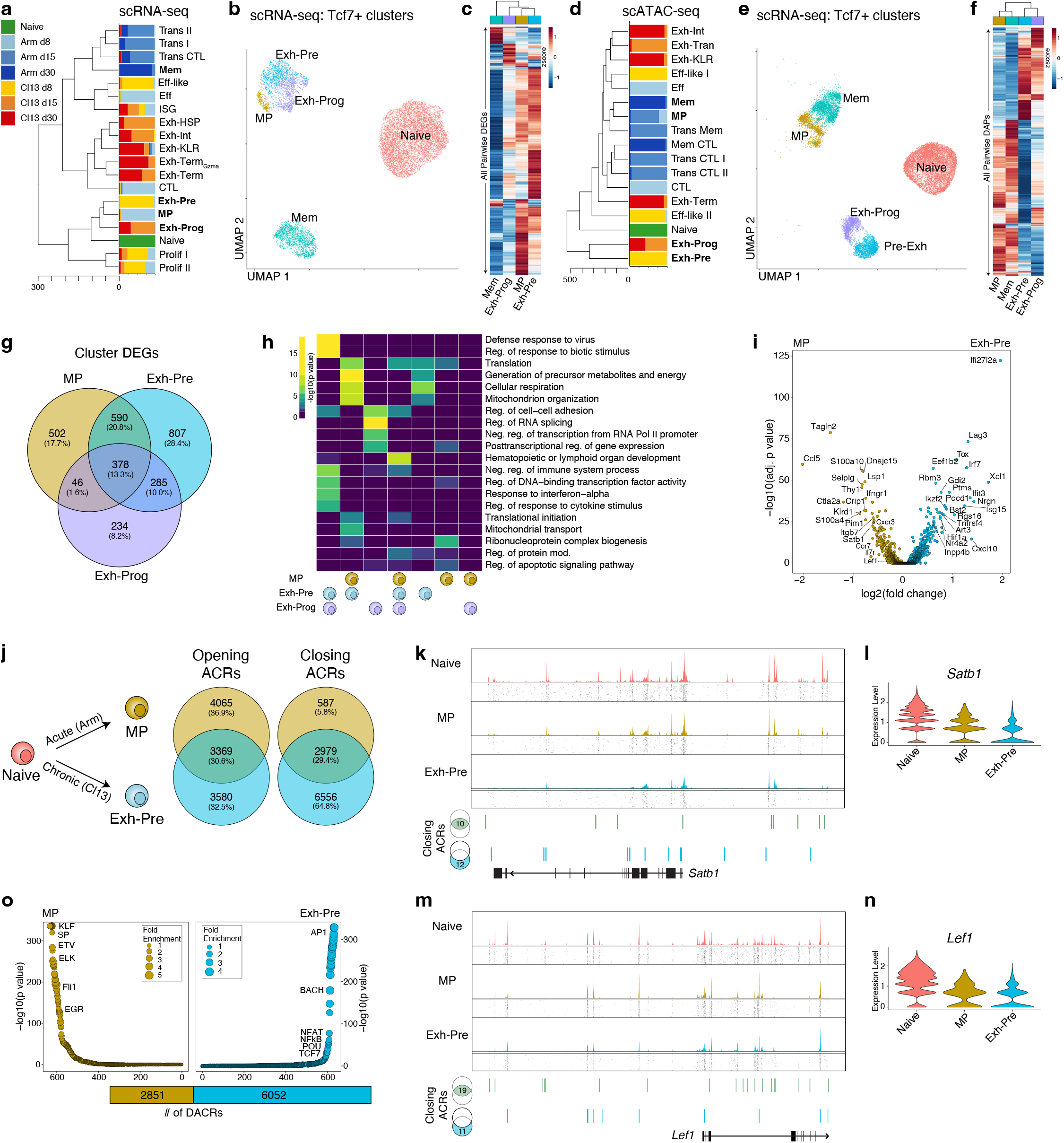
Acute and chronic infections generate *Tcf7*-expressing progenitors with divergent accessible chromatin profiles. Phylogenetic tree of (**a**) scRNA-seq or (**d**) scATAC-seq clusters; bar plots represent proportion of cells per infection and timepoint. UMAP from (**b**) scRNA-seq or (**e**) ATAC-seq of Naïve, MP, Mem, Exh-Pre, Exh-Prog subsets, colored by subset. **c**) Average gene expression per cluster of all pairwise DEGs. **f**) Average accessibility per cluster of all pairwise DACRs. **g**) Venn diagram of MP, Exh-Pre, Exh-Prog DEGs. **h**) Gene ontology of DEGs in (**g**). **i**) DEGs between scRNA-seq MP and Exh-Pre clusters. **j**) Venn diagram of DACRs calculated between scATAC-seq naïve and MP or naïve and Exh-Pre clusters. Coverage and tile plots from scATAC-seq of *Satb1* locus (**k**) or *Lef1* locus (**m**); DACRs closing in both MP and Exh-Pre (green) or only in Exh-Pre (blue), compared to Naïve, indicated on the bottom. Gene expression of *Satb1* (**l**) or *Lef1* (**n**). **o**) TF motif enrichment of DACRs comparing MP and Exh-Pre; total number of DACRs represented on bottom.

The results above revealed transcriptional similarity among MP, Exh-Pre, and Exh-Prog subsets. This observation may reflect a convergence of gene expression related to cell activation. We investigated this possibility by comparing the gene expression profiles of these subsets. MP and Exh-Pre shared expression of 968 genes, 378 of which were also expressed by Exh-Prog (**Fig. 6g**). Among these three cell types, the Exh-Pre cluster had the greatest number (807) of uniquely expressed genes (**Fig. 6g**). Gene ontology analysis revealed that many pathways shared between MP and Exh-Pre were related to cellular metabolism, including cellular respiration, generation of metabolites, and mitochondrial function (**Fig. 6h**), consistent with the notion that *Tcf7+* cells, despite their “stem-like” features, are also in a highly active state early during both acutely-resolving and chronic infection. However, the expression of viral-response gene programs in Exh-Pre and Exh-Prog, but not MP, point to induction of some key distinct pathways in Cl13 at d8 compared to Arm. Despite some shared pathways, MP and Exh-Pre differed in expression of the exhaustion-driving TF *Tox*, consistent with previous data (22), as well as the IR *Lag3* and many ISGs (e.g. *Ifi27I2a, Isg15, Ifit3*, and *Irf7*) (**Fig. 6i**). Thus, MP and Exh-Pre subsets in acute and chronic infection, share transcriptional features of T cell activation and metabolic activity that may drive co-localization in scRNA-seq space. Nevertheless, Exh-Pre have a distinct transcriptional program that includes key exhaustion-specific TFs and IRs.

Given the epigenetic divergence in subsets from acute versus chronic infection, we next directly compared chromatin accessibility changes between Naïve and d8 precursor cells in Arm (MP) versus Cl13 (Exh-Pre) (**Fig. 6j**). Among regions that increased in accessibility, one third were shared and one third each were unique to MP or Exh-Pre. In contrast, most DACRs that lost accessibility were unique to Exh-Pre (6556 ACRs, ~65%). MP only had 587 regions that closed (~6%), and 2979 ACRs (~30%) were closed in both MP and Exh-Pre. Some regions that lost accessibility between Naïve and Exh-Pre cells were near genes related to self-renewal, including *Sat1b* (41) and *Lef1* (42). At the *Satb1* locus, 10 ACRs lost accessibility in both MP and Exh-Pre; however, an additional 12 were closed only in Exh-Pre (**Fig. 6k**), and this pattern was mirrored in the gene expression profiles (**Fig 6l**). *Lef1* followed a similar pattern (**Fig. 6m,n**). Thus, one major distinction of T_EX_ progenitors in chronic infection may be decreased expression of stem-associated genes, a set of changes that may prevent full conversion to quiescence. Finally, we directly compared ACRs in Exh-Pre and MP. This analysis revealed enrichment of AP1 TF binding motifs in Exh-Pre-specific ACRs (**Fig. 6o**), suggesting a prominent role for TCR signaling in shaping the Pre-Exh epigenetic landscape and/or TCR-dependent TFs operating in this ACR landscape. In contrast, MP were enriched in accessibility for ETS family TFs, including motifs for Fli1, a TF that may help restrain CD8 T cell activation (43). Altogether, these data reveal distinct paths of *Tcf7*-expressing cells early during acutely-resolved versus chronic infection and identify different biological modules that can be present in TCF1+ “stem” or “progenitor”-like CD8 T cells.

### Biological circuits underlying the transition from Exh-Pre to Exh-Prog

Finally, we investigated transcriptional and epigenetic changes between Exh-Pre and Exh-Prog because this transition marks a point of irreversibility in commitment to exhaustion (16, 39, 44). We directly compared transcriptional profiles of Exh-Pre and Exh-Prog. Almost 1000 genes were increased in Exh-Pre from d8 compared to the Exh-Prog from d15, but there were few transcriptional differences between Exh-Prog at d15 and Exh-Prog at d30 (**Fig. 7a**), consistent with establishment of T_EX_ by d15 and stabilization of the Exh-Prog transcriptional program. Exh-Pre DEGs were enriched in pathways related to metabolism and mitochondrial function (**Fig. 7b**), supporting the results above indicating Exh-Pre cells are highly activated at d8 p.i.. Here, we found a decrease in these pathways from Exh-Pre to Exh-Prog as well as a decrease in protein translation (**Fig. 7b**). Because protein translation is one of the most bioenergetically costly cellular activities (45), it may be challenging to sustain at a high level over time in T_EX_ despite ongoing activation signals from antigen stimulation. We used an *in vitro* translation assay that measures uptake of a fluorescently labeled amino acid analog, l-homopropargylglycine (HPG) to assess changes in protein translation from d8 to d15. At d8, Exh-Pre from Cl13 had significantly higher HPG incorporation than MP from Arm (**Fig. 7c**). However, by d15 in Cl13, this HPG signal was substantially reduced in Exh-Prog (**Fig. 7c**). These data indicate that despite ongoing antigen stimulation during chronic infection, one major feature of the Exh-Pre to Exh-Prog transition is dampening metabolic and protein translation activities. Establishing a more quiescent state juxtaposed to strong continued stimulation may be necessary to ensure cellular persistence in chronic infection. In contrast to the scRNA-seq data that indicated increased transcriptional activity in Exh-Pre, scATAC-seq revealed a greater number of ACRs in d15 Exh-Prog compared to d8 Exh-Pre (**Fig. 7d**). Several gene loci had multiple DACRs including *Fos, Fosb, Dusp1, Tnfaip3* and *Btg1* (**Fig. 7e**). AP1 family member genes were notable given their importance in exhaustion and effector biology (46–49). *Tnfaip3* is known to regulate CD8 T cell effector functions through inhibition of NFkB (50, 51). *Btg1* was of particular interest because of its role in maintaining homeostasis under stress (52) including in hematopoietic stem cells returning to quiescence after proliferation (53).

**Figure 7.**
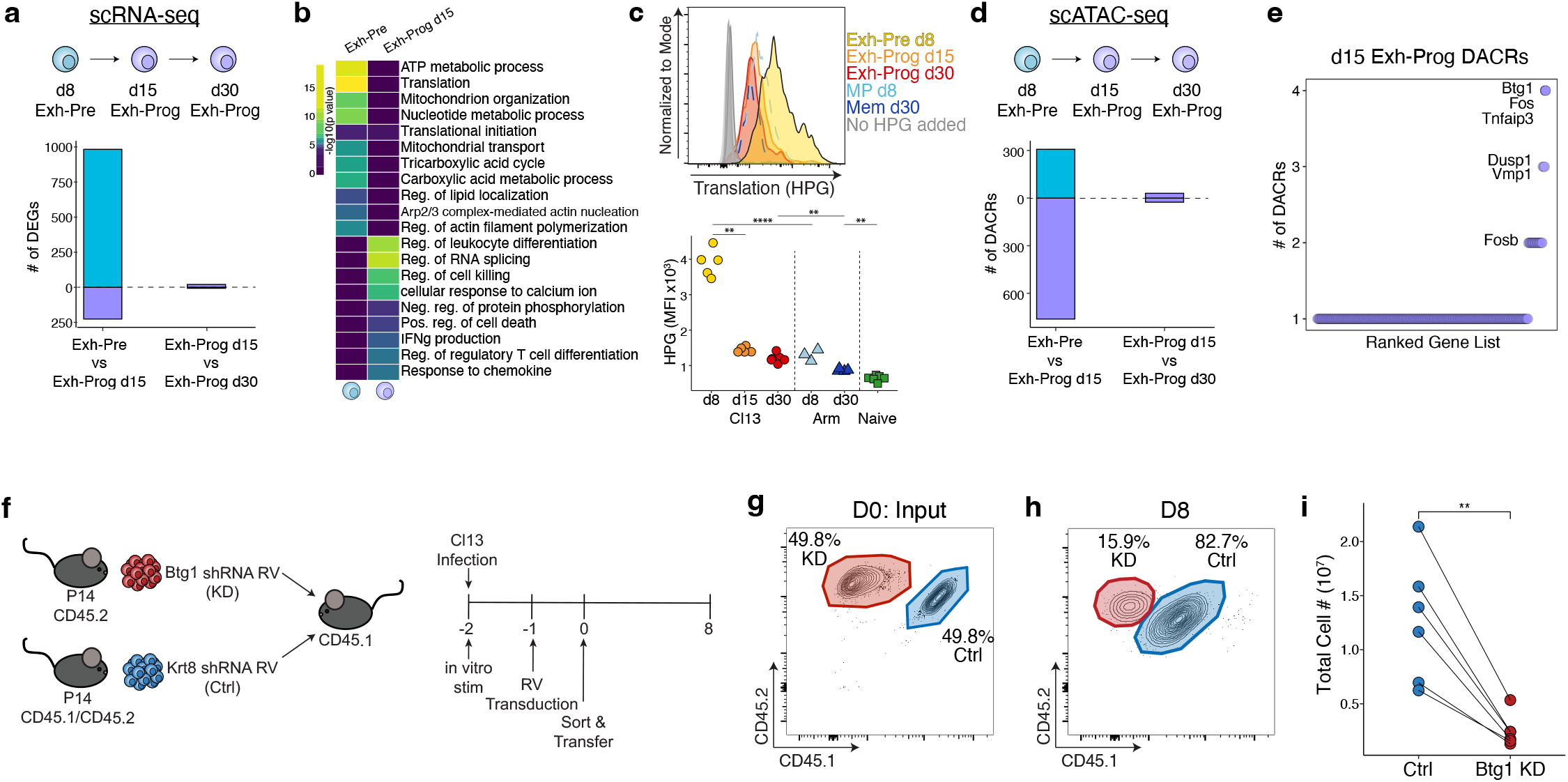
Transition from Exh-Pre to Exh-Prog uncovers Btg1 as a novel regulator of T_EX_ differentiation. **a**) Enumerated DEGs from pairwise analysis as indicated. **b**) Gene ontology of DEGs from (**a**) as indicated. **c**) Representative flow cytometry plot (top) and enumeration of MFI of HPG signal (bottom) from *in vitro* translation assay. Cells are gated as described in **Figure 3j**. Data are representative of two experiments with 3-5 mice per group. **d**) Enumerated DACRs from pairwise analysis as indicated. **e**) For DACRs increased in d15 Exh-Prog, number of DACRs per gene loci, select genes overlapping with DEGs annotated. **f**) Experimental schematic. **g**) Flow cytometry plot of input P14 cell mixture containing *Btg1* KD and *Krt8* Ctrl. **h**) Representative flow cytometry plot from d8 p.i. Cells are gated on RV^+^ (GFP) live P14 T cells. Mean percentage as indicated. **i**) Total P14 cells with Ctrl or *Btg1* KD RV. Comparison with two-sided t-test ** p<0.01. Representative data are from two experiments. Each point represents a mouse.

Lastly, to test whether Btg1 had a role in T_EX_ differentiation, we used retroviral (RV)-mediated shRNA knockdown (KD) (**Fig. 7f** and **Fig. S5**). We transduced LCMV-specific P14 cells with RV encoding shRNA targeting *Btg1* or *Krt8* (an irrelevant control gene; Ctrl) followed by dual adoptive transfer into congenically distinct mice infected with Cl13 (**Fig. 7f**). Despite an equal mixture of cells targeting the Ctrl versus *Btg1* in the input population (**Fig. 7g**), *Btg1* KD resulted in significantly fewer T_EX_ by d8 (**Fig. 7h-i**). The analyses above suggest a role for *Btg1* in regulating the transition from the highly stimulated Exh-Pre population present in the first week of chronic infection to a more “regulated” Exh-Prog population the emerges by d15. Here, we find that KD of *Btg1* had a profound effect early, by d8 after infection. Together, these data indicate a key role for this stress response gene in the ability to generate early T_EX_ cells and in the transition from the early phase of exhaustion to formation of established T_EX_. These data also suggest that *Btg1* and related cellular stress response genes could be attractive therapeutic targets to modulate T_EX_ populations.

## Discussion

We used the LCMV mouse model of CD8 T differentiation to generate T_EFF_, T_MEM_, and T_EX_, and performed scRNA-seq and scATAC-seq over time to investigate the transcriptionally- and epigenetically-defined subset heterogeneity and underlying molecular development pathways. These analyses revealed several key insights. First, scATAC-seq defined fewer clusters compared to scRNA-seq, demonstrating that multiple transcriptional states can exist from a smaller number of epigenetic cell fates. Gene expression might have less resolution in defining cell identity due to convergent patterns of gene expression (e.g. activation or cell cycle) from distinct cell types. This data supports the idea that chromatin accessibility profiles are better suited to define cell “fates”. Second, these data uncovered new subpopulations of T_EFF_, T_MEM_, and T_EX_, including a T_EX_ subset expressing NK receptors (Exh-KLR) and an early T_MEM_ subset distinguished by cytolytic potential (Mem-CTL). Third, analyzing chromatin accessibility after PD1 pathway blockade revealed a shift in T_EX_ subsets including preferential expansion of the Exh-Int subset at the expense of the Exh-KLR subset and evidence of repopulating more terminal T_EX_ subsets with new cells. Fourth, we identified multiple epigenetically distinct TCF1+ CD8 T cell populations in chronic and acute-resolving infection. Although some of these TCF1+ populations had shared transcriptional features, they were imprinted with unique accessible chromatin landscapes that further evolved over time as T_MEM_ and T_EX_ developed. Lastly, we identified the stress response gene, *Btg1*, as a novel regulator of CD8 T cell exhaustion. Altogether, this single cell transcriptional and chromatin accessibility landscape map of CD8 T cell differentiation using a defined system provides insights into the developmental biology and underlying mechanisms governing T_EFF_, T_MEM_, and T_EX_ formation and regulation.

One major observation from this study was identification of a subpopulation of T_EX_ cells expressing NK-associated genes. Recent work has highlighted NK receptor-expressing CD8 T cells in cancer as potentially therapeutically relevant (32, 34, 35, 54). It has been unclear, however, whether these CD8 T cells represent a distinct subset or reflect transient expression of NK-associated genes (e.g. akin to an activation event). Here, we found that CD8 T cells expressing NK-associated genes form a distinct cluster of T_EX_ in unbiased analysis of scRNA-seq data. Moreover, this Exh-KLR subset was readily apparent by scATAC-seq, pointing to an underlying chromatin-accessibility driven cell fate rather than a transient transcriptional state. A key question emerging from these observations is whether this subset only arises in chronic infection. Because we examined both acute-resolving and chronic infection, we were able to identify populations of CD8 T cells from both Arm and Cl13 infection that expressed NK receptors, including NKG2A and CD94, consistent with early studies documenting expression of NK receptors on CD8 T cells in viral and bacterial infections (37, 38). It was clear that although these NK-receptor expressing CD8 T cell subsets in Arm and Cl13 infection shared this biological circuit, these subsets were otherwise largely distinct cell types. Future work will be needed to determine the functional significance of NK receptor expression on CD8 T cells but identification of a discrete Exh-KLR subset within the T_EX_ population suggests these cells may play a unique role in sensing infected or stressed cells through these NK receptors.

A second key finding from these studies was identification of multiple epigenetically distinct TCF1+ stem-like CD8 T cell populations in different disease settings across time. “Stem-like” or progenitor CD8 T cells in chronic infection and cancer have received considerable recent attention given their role in response to immune checkpoint blockade (7, 9, 14, 55, 56) and adoptive T cell therapy (10). However, it has been unclear whether all stem/progenitor-like CD8 T cells are the same or whether different types of TCF1+ CD8 T cells exist in different settings. Our data now clearly demonstrate the existence of multiple epigenetically distinct populations of TCF1+ stem/progenitor-like CD8 T cells in acutely-resolved and chronic viral infection. Therefore, TCF1 expression alone in non-naïve CD8 T cells is not sufficient to define the biology of these stem/progenitor populations. Specifically, these data show that at d8 p.i. MP and Exh-Pre form distinct cell types defined by either transcriptional or epigenetic information. Gene expression patterns identified both MP and Exh-Pre as highly activated early after infection revealing a point of biological convergence. However, other key biology was markedly distinct including substantially higher expression of the master regulator of T_EX_ epigenetics TOX in Exh-Pre consistent with previous studies (22). Although Exh-Pre and MP share transcription features of activation, chromatin accessibility patterns were highly divergent with Exh-Pre sharing major features of the T_EX_ epigenetic landscape even early after activation, consistent with high TOX expression, whereas MP preserved more stem-like biology. Previous studies indicated that full commitment to exhaustion occurs progressively over time (18, 39) and becomes largely irreversible by ~two weeks (16, 44). Indeed, Exh-Prog, first identifiable at d15, were transcriptionally and epigenetically distinct from these early Exh-Pre cells. Transitioning from Exh-Pre to Exh-Prog was accompanied by dampening many cellular processes, including metabolism and translation. This relative return to quiescence may be a key event that allows the T_EX_ population to persist despite strong ongoing antigenic stimulation. The ability to distinguish between Exh-Pre and Exh-Prog may be particularly relevant in settings where initial activation is not synchronized. For example, tumor-specific T cells in cancer may get primed at different points as cancer cells mutate and evolve over time. This asynchronous activation could be relevant because recently activated Exh-Pre retain more fate flexibility (44) and would be predicted to respond differently than Exh-Prog to immunotherapies. Disentangling closely related but distinct CD8 T cell populations such as Exh-Pre and Exh-Prog could have key relevance for understanding immune responses after treatment and for identifying clinical biomarkers.

By analyzing transcriptional and epigenetic networks over time, we identified TFs involved in T_EX_ biology. The stress regulator, *Btg1*, was necessary for sustaining Exh-Pre and Eff-like cells early in chronic infection and was predicted to regulate the transition from highly activated Exh-Pre to established T_EX_ populations. In hematopoietic stem cells, *Btg1* regulates quiescence and survival after proliferation (52, 53). In T_EX_, the requirement of *Btg1* likely reflects a similar role in attenuating robust cellular activation despite ongoing TCR signaling from persistent antigen stimulation. *Btg1* and related targets may provide novel opportunities to foster (in the case of cancer or chronic infections) or eliminate (in the case of autoimmunity) more durable T_EX_ populations.

In summary, scRNA-seq and scATAC-seq landscapes of T_EFF_, T_MEM_, and T_EX_ revealed subpopulation heterogeneity and developmental trajectories. Comparative analysis across these cell types identified shared and distinct transcriptional and epigenetic programs underlying cellular identities. These data overall highlight a key theme of “re-using” biological circuits in different CD8 T cell populations. This concept was apparent for NK-associated cytotoxicity and TCF1-centered progenitor biology that were both found in epigenetically distinct CD8 T cell subpopulations. Thus, this transcriptional and chromatin accessibility landscape map provides insights into the developmental biology and underlying mechanisms governing T_EFF_, T_MEM_, and T_EX_ differentiation and may help identify specific targets or pathways for future therapeutic manipulation.

## Acknowledgements

We thank members of the Wherry Lab. This work was supported by T32 CA009140 and a Cancer Research Institute-Mark Foundation Fellowship (JG), by the Parker Institute for Cancer Immunotherapy and Stand Up To Cancer and NIH grants AI155577, AI115712, AI117950, AI108545, AI082630 and CA210944 (to EJW). Work in the Wherry lab is supported by the Parker Institute for Cancer Immunotherapy. SFN was supported by an Australia NHMRC C.J. Martin Fellowship (GNT1111469) and the Mark Foundation Momentum Fellowship. OK was supported by an NIAID F30 fellowship (F30AI129263). DM was supported through The American Association of Immunologists Intersect Fellowship Program for Computational Scientists and Immunologists. JEW was supported by a PICI Scholar award. YJH was supported by a National Science Foundation graduate research fellowship.

## Author Contributions

JG, OK, and EJW conceived and designed the experiments. JG, OK, RS performed FACS and prepared sequencing libraries. JG analyzed data with help from SFN, SM, and HH. PW prepared retroviruses. MSA provided long-term Arm infected mice. AEB, SFN, DM, MMP, RG, JEW, YJH helped with experiments. JG and EJW wrote the manuscript.

## Supplemental Figure Legends

**Figure S1.**
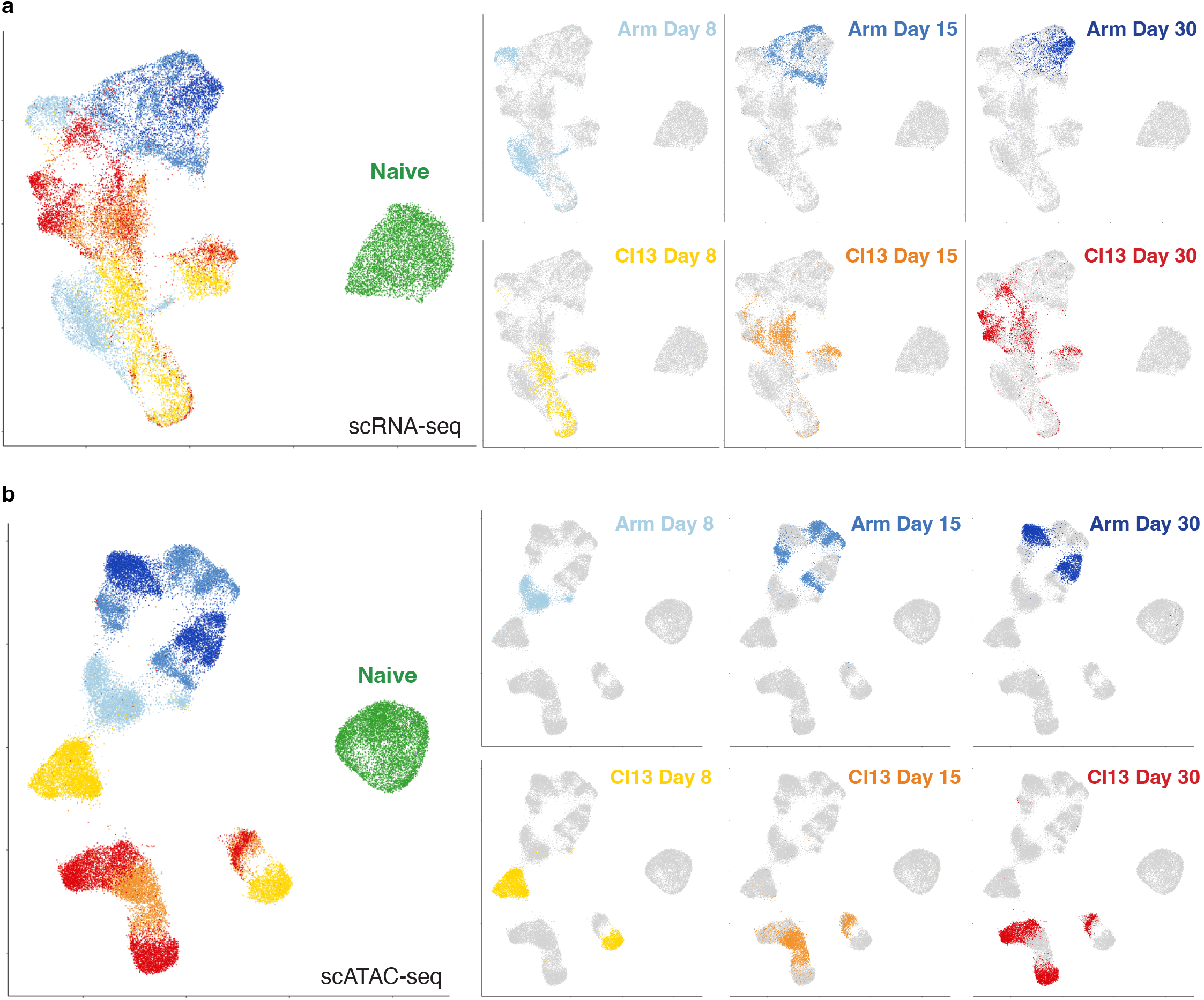
UMAP analysis of scRNA-seq and scATAC-seq by infection and timepoint. UMAP from (**a**) scRNA-seq and (**b**) scATAC-seq colored by infection and timepoint as indicated.

**Figure S2.**
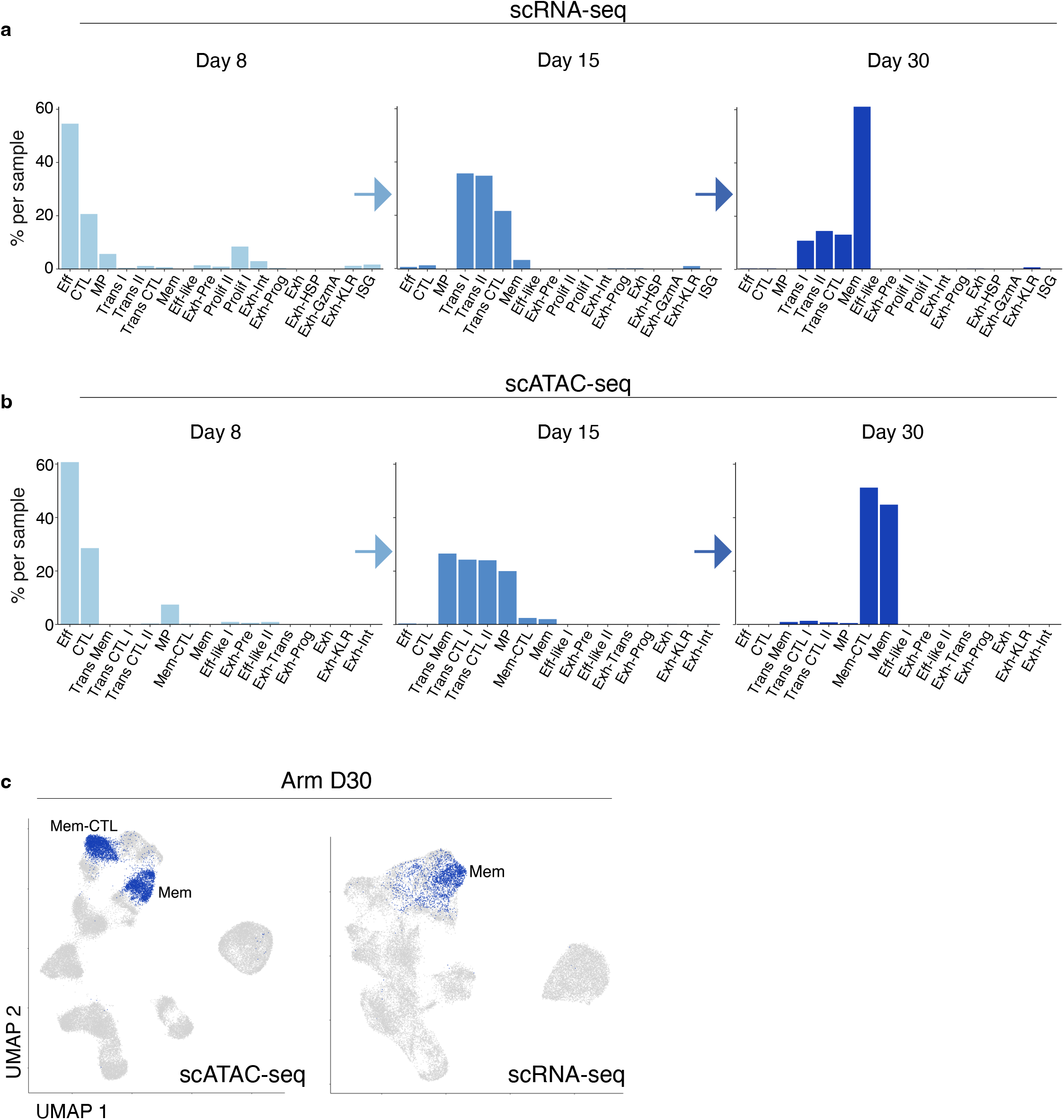
Effector and memory clusters defined by scRNA-seq and scATAC-seq data identify shared and non-overlapping cell subsets. Percentage of cells from Arm infection by timepoint as indicated in (**a**) scRNA-seq clusters and (**b**) scATAC-seq clusters. **c**) scATAC-seq UMAP (left) and scRNA-seq UMAP (right) colored with d30 Arm cells.

**Figure S3.**
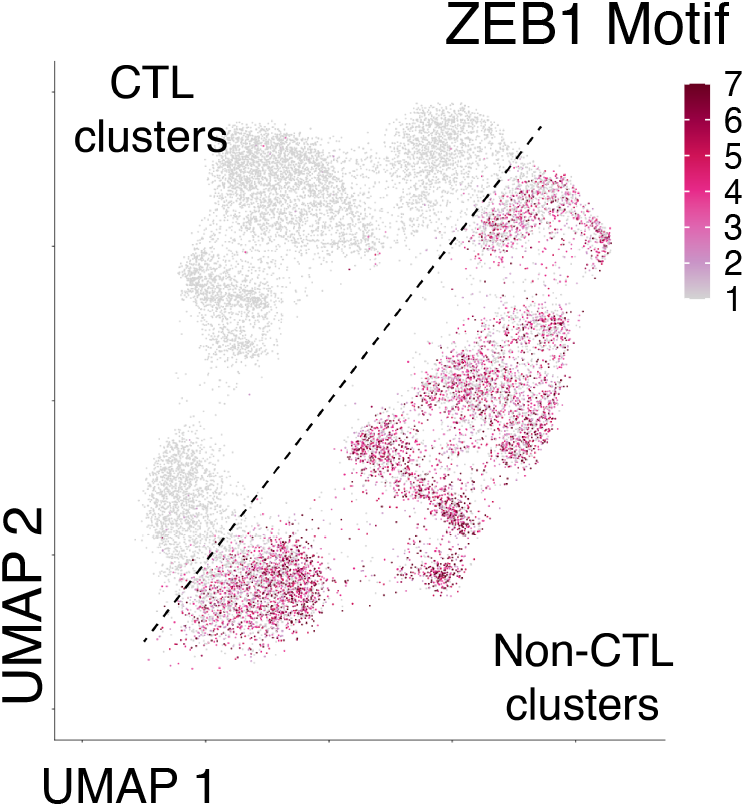
ZEB1 motif is enriched in non-CTL clusters. scATAC-seq UMAP of cells from Arm infection colored by ZEB1 motif enrichment. The location of CTL and non-CTL clusters is indicated.

**Figure S4.**
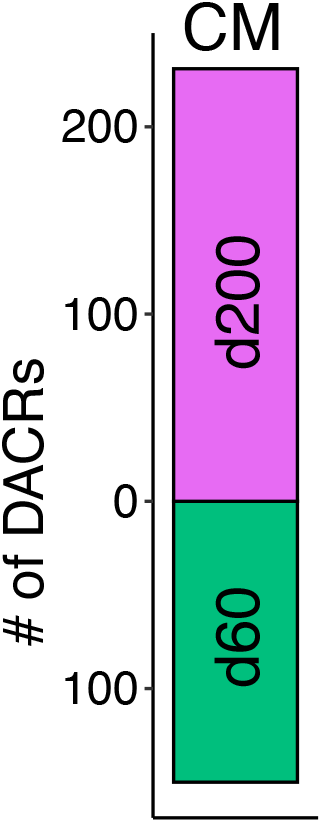
The accessible chromatin landscape in central memory CD8 T cells evolves over time. Number of DACRs between CM d60 and CM d200 clusters.

**Figure S5.**
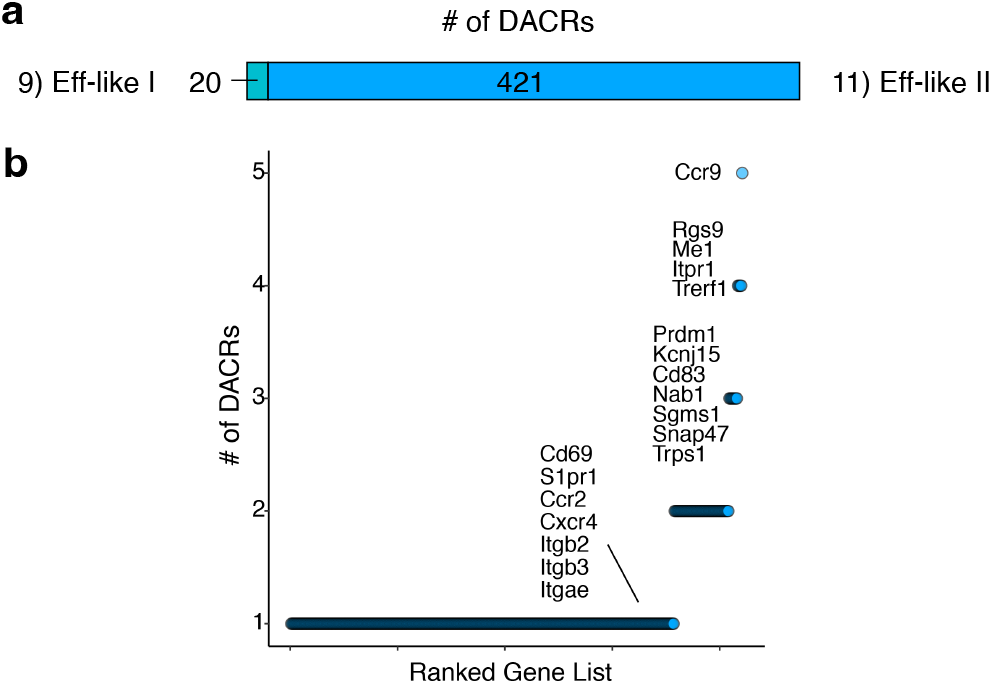
scATAC-seq defined clusters Eff-like I and Eff-like II are distinguished by DACRs at gene loci related to migration. **a**) Barplot representing the number of DACRs between scATAC-seq clusters Eff-like I and Eff-like II. **b**) Number of Eff-like II DACRs per gene loci. Genes of interest manually annotated.

**Figure S6.**
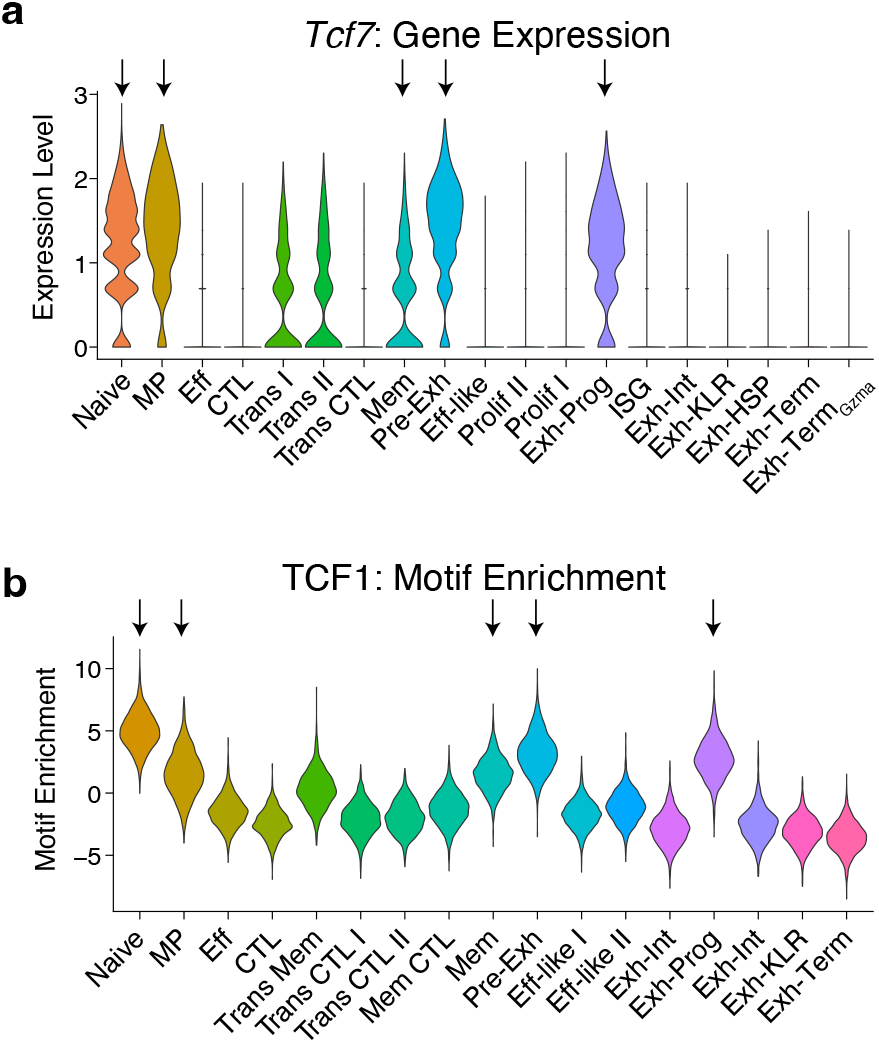
Identification of Tcf7-expressing progenitor/stem-like CD8 T cell subsets. **a**) Gene expression from scRNA-seq of all scRNA-seq defined clusters. **b**) Motif enrichment from scATAC-seq of all scATAC-seq defined clusters.

**Figure S7.**
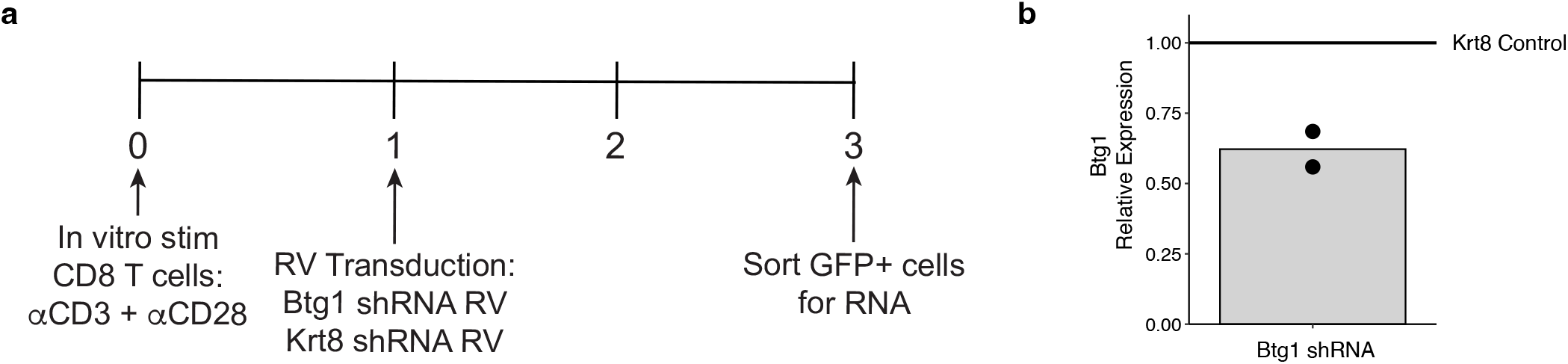
Retroviral-mediated knock down of Btg1. **a**) Experimental schematic. **b**) qPCR results of shRNA-mediated knockdown of Btg1. Bar represents mean, points represent independent experiments.

## Methods

### Data availability

All scRNA-seq and scATAC-seq data generated in this study will be deposited in GEO by time of publication.

### Code availability

All analyses were done with custom R scripts that can be made available upon request using standard R packages. No new algorithms were developed during this study.

### Mice

P14 transgenic mice expressing a TCR specific for the LCMV peptide gp33-41 were bred in-house and backcrossed onto the C57BL/6 background. Recipient mice were purchased from Jackson Laboratory or Charles River. All experiments and breeding conditions were in accordance with Institutional Animal Care and Use Committee (IACUC) guidelines for the University of Pennsylvania.

### Adoptive T cell transfer

Recipient mice were adoptively transferred with PBMCs containing P14 CD8 T cells isolated from peripheral blood of donor P14 mice using gradient centrifugation with Histopaque-1083 (Sigma-Aldrich). For most experiments, 500 naïve P14 cells were adoptively transferred intravenously (i.v) into 6-7-week-old sex-matched recipient mice 1 day prior to infection. In long-term Arm experiments (d60 and d200), 5,000-10,000 naïve P14 cells were transferred to facilitate adequate cell recovery at late time points after infection. Recipients were of a distinct congenic background to allow for identification of donor populations from host CD8 T cells.

### Infections

LCMV Armstrong and clone 13 were grown in BHK cells and titrated using plaque assay on VERO cells. Recipient mice were infected intraperitoneally (i.p.) with LCMV Armstrong (2×10^5^) plaque forming units (PFU) or i.v. with LCMV clone 13 (4×10^6^ PFU) one day post adoptive transfer of P14 cells. The number of mice infected per condition was 10 for d8 Arm, 15 for d15 Arm, 15 for d30 Arm, 10 for d8 Cl13, 20 for d15 Cl13, 20 for d30 Cl13, 15 for d30 Cl13 + αPDL1. For the long-term Arm memory experiment, 4 mice were infected for each d60 and d200.

### PD1 blockade

PD1 blockade was performed with 5 treatments of 200ug aPD-L1 antibody (10F.9G2, BioXCell) i.p. every 3 days starting 16 days after infection with LCMV Cl13. Analysis was performed one day after final treatment. For control treatments, phosphate-buffered saline (PBS) was administered i.p. The blockade experiment was performed at the same time as the experiment in **Fig. 1a**.

### Cell sorting for sequencing libraries

Spleens from mice in the same experimental group (i.e. d8 Arm, etc.) were processed together, five at a time. Spleens were homogenized using a Miltenyi gentleMACS™ Dissociator in C tubes. CD8 T cells were enriched using an EasySep magnetic negative selection kit (Stem Cell Technologies #19853) as per the manufacturer’s recommendations. Cells were washed with 1× PBS and stained with an amine–reactive dye (Biolegend # 423106) for 20 minutes at room temperature to assess cell viability, followed by an antibody cocktail in complete RPMI (cR10, RPMI-1640 media supplemented with 10% FBS, 1x non-essential amino acids (Gibco #11140050), and 10mM Hepes (Gibco #15630080), 2mM L-glutamine (Gibco #25030081), 100U/mL penicillin/streptomycin (Gibco #15140122), 14.3 uM beta-mercaptoethanol) for 45 minutes on ice. Samples were sorted on a BD FACSAria II machine into RPMI-1640 media supplemented with 50% FBS, 1% Hepes, 1% L-glutamine, 1% penicillin/streptomycin (CR50). Cells gated as live single CD8+ P14 cells designated by congenic markers. A small aliquot of all sorted samples was run as a purity check. Voltages on the machine were standardized using fluorescent targets and Spherotech rainbow beads (#URCP-50-2F).

### Flow cytometry

Single cell suspensions were prepared by mechanically disrupting spleen through a 70 μM cell strainer using the plunger of a 3mL syringe; followed by red blood cell lysis with ACK buffer (Gibco #A10492-01). Cells were washed with PBS and stained with an amine–reactive dye (Biolegend #423104) for 20 minutes to assess cell viability. Surface staining was performed for 45 minutes at RT in staining media (SM), PBS with 3% FCS, 5mM EDTA, and 1% penicillin/streptomycin, followed by secondary stain using streptavidin-Brilliant Blue 790 (BD Biosciences) in SM for 30 minutes on ice. Permeabilization was performed using the Foxp3 Fixation/Permeabilization Concentrate and Diluent kit (eBioscience #00-5521-00) for 20 minutes. Intracellular staining with antibody cocktails was done for 2 hours at room temperature. Samples were run on a BD FACSymphony A5 instrument. Voltages on the machine were standardized using fluorescent targets and Spherotech rainbow beads (#URCP-50-2F). Data were analyzed with FlowJo software (version 10.5.3, TreeStar).

### Translation Assay

The protein translation assay was adapted from (58). First, single cell suspensions were prepared as described above (see “Flow cytometry” section). Then, the cells were washed and plated at 1 million cells per well in a V-bottom 96 well plate in methionine-free R10 (Gibco #A1451701) supplemented with 10% FBS, 1x non-essential amino acids (Gibco #11140050), 10mM Hepes (Gibco #15630080), 2mM L-glutamine (Gibco #25030081), 100U/mL penicillin/streptomycin (Gibco # 15140122), 14.3uM beta-mercaptoethanol. The cells were rested at 37°C for 3 hours, then 400uM Click-iT HPG (Invitrogen #C10186) was added. After 3 hours, cells were stained with viability dye and surface antibody cocktail as described above (see “Flow cytometry” section). Then, the cells were fixed and permeabilized (BD #51-2090KZ) for 20 minutes at RT, followed by one wash with perm wash (BD #51-2091KZ) and one wash with PBS. Next, the Click-It reaction was performed per manufacture’s protocol (Invitrogen #C10641). Samples were analyzed as described above (see “Flow cytometry” section).

### shRNA Design and Retroviral Transduction

shRNA sequences and cloning strategy are as described in (59). Briefly, 97-mer shRNA oligonucleotides were synthesized (IDT) and 4 picomoles were amplified with HotStarTaq polymerase (Qiagen#203207) using the primers miR-E-fw (5’-TGAACTCGAGAAGGTATATTGCTGTTGACAGTGAGCG-3’) and miR-E-rev (5’-TCTCGAATTCTAGCCCCTTGAAGTCCGAGGCAGTAGGC-3’). Following amplification, reactions were purified (Qiagen MinElute PCR Purification Kit; #28004) and subsequently digested with XhoI/EcoRI using standard techniques. Amplicons were purified (Qiagen MinElute Reaction Cleanup Kit; #28206) and ligated into XhoI/EcoRI-digested LMPd plasmid (kindly provided by the Shane Crotty laboratory) with T4 DNA ligase. Sequence-verified plasmids were then used to transform TOP10 chemically competent bacterial cells (ThermoFisher; #C404010) and endotoxin-free plasmid stocks were prepared (Qiagen EndoFree Plasmid Maxi Kit; #12362). RV was generated for each construct as previously described (60).

For RV transduction, single cell suspensions were prepared by mechanically disrupting spleen through a 70 μM cell strainer using the plunger of a 3mL syringe. CD8 T cells were enriched using an EasySep magnetic negative selection kit (Stem Cell Technologies) as per the manufacturer’s recommendations. P14 T cells were stimulated with αCD3 (1mg/mL), αCD28 (0.5mg/mL), and IL-2 (100U/mL) (PeproTech). Thirty hours post activation, T cells were spin-transduced for 75 minutes at 2000g 37°C with RV supernatant containing polybrene (4mg/mL) and IL-2 (100U/mL). Approximately 24 hours later, GFP positive cells were sorted on a BD FACSAria II machine into cR50. A small aliquot of all sorted samples was run as a purity check. Voltages on the machine were standardized using fluorescent targets and Spherotech rainbow beads (#URCP-50-2F). Sorted cells were washed twice with warm unsupplemented RPMI. An equal number of cells transduced with the control RV (*Krt8*) or experimental RV (*Btg1*) (25,000 *Krt8* Ctrl + 25,000 *Btg1*) were transferred i.v. into mice that had been infected with LCMV Cl13 two days before (the same day as *in vitro* stimulation).

### scRNA-seq library generation

scRNA-seq libraries were generated using the 10x Genomics Chromium Single Cell 3’ Library (v2). In brief, sorted P14 T cells were washed with 0.04% BSA PBS, then approximately 20,000 cells were loaded into a 10x Chromium controller. All downstream library preparation steps were performed according to manufacturer’s instructions. Libraries were assessed using an Agilent Tapestation and quantified using a KAPA Library Quantification Kit (#KK4824) and sequenced on an Illumina Novaseq.

### scATAC-seq library generation

scATAC-seq libraries were generated using the 10x Genomics Chromium Cell ATAC Reagent Kit (v1). In brief, sorted P14 T cells were washed with 0.04% BSA PBS, then approximately 40,000 cells were subjected to the nuclei preparation protocol per manufacturer’s instructions. Then, 16,000 nuclei were loaded into a 10x Chromium controller. All downstream library preparation steps were performed according to manufacturer’s instructions. Libraries were assessed using an Agilent Tapestation and quantified using a KAPA Library Quantification Kit (#KK4824) and sequenced on an Illumina Novaseq.

### scRNA-seq data processing and analysis

scRNA-seq data was generated using the 10x Cell Ranger pipeline (3.0.2) and mm10 genome. Specifically, we generated fastqs using cellranger mkfastq, then quantified reads using cellranger count, and cellranger aggr to combine samples. Downstream analysis was performed in R (version 4.0.2) and Seurat (version 4.0.4) using default parameters unless otherwise noted. Cells with less than 200 features and more than 0.75% mitochondrial reads were excluded. Standard Seurat data processing and normalization steps were performed: SCTransform, RunPCA, RunUMAP, FindNeighbors, and FindClusters; clusters with low quality scores were removed, final resolution 0.9. Clusters 9 and 13 which split in a high resolution were kept separate based on their large number of DEGs. Proliferation analysis used the CellCycleScoring function (Seurat). DEGs were calculated using the functions, FindAllMarkers (Seurat) or FindMarkers (Seurat), for pairwise comparisons using a log2 fold change threshold of 0.125, adjusted p value less than 0.05, and include the number of counts as a latent variable. Gene set enrichment was performed using the AddModuleScore function (Seurat), and gene ontology analysis of DEGs used metascape (metascape.org) with all expressed genes as the background gene list. Phylogenetic trees were constructed with the BuildClusterTree function (Seurat). Average gene expression was calculated using the AverageExpression function (Seurat). All heatmaps were generated using pheatmap (version 1.0.12). Cluster prediction of proliferating cells (**Fig. 3g**) was accomplished by creating two Seurat objects, one with proliferating cells and one without proliferating cells using the standard processing steps described above and including S.Score and G2M.Score calculated from CellCycleScoring as variables to regress in the SCTransform calculation. The proliferating cells were then projected onto the UMAP of non-proliferating cells using the Seurat functions FindTransferAnchors, TransferData, IntegrateEmbeddings, and ProjectUMAP.

### scATAC-seq data processing and analysis

scATAC-seq data was generated using the 10x Cell Ranger ARC pipeline (2.0.0) and mm10 genome. Specifically, we generated fastqs using cellranger mkfastq, then quantified reads using cellranger arc-count. Downstream analysis was performed in R (version 4.0.2), Seurat (version 4.0.4), Signac (1.3.0), and ArchR (1.0.1) using default parameters unless otherwise noted. ArchR was used to perform initial quality control (TSSEnrichment > 10, nFrag > 1500 & < 30000, BlacklistRatio < 0.1) and identify doublets. The union peak list was generated using a hybrid approach. Peaks were called using ArchR with default parameters based on clusters generated from the LSI dimension reduction of the tile matrix which allowed peaks to be called on unbiased cell clusters. In addition, we called peaks on the sample bam files (naïve, Arm d8, Arm d15, Arm d30, Cl13 d8, Cl13 d15, Cl13 d30, Cl13 d30+αPDL1) using macs2 as previously described (23) with a q value 0.001, then combined the two peak lists. Downstream analysis was performed using the Signac package, unless otherwise noted. In addition to the ArchR metrics, quality control metrics were also calculated in Signac and cells were filtered as nCounts_peaks > 3500 & < 35000, blacklist_fraction < 0.035, nucleosome_signal < 5. The custom peak list was added to the Signac object using FeatureMatrix and CreateChromatinAssay. Peak annotation was performed using the ClosestFeature function. Standard processing and normalization steps were performed: FindTopFeatures, RunTFIDF, RunSVD, and FindClusters (resolution 0.9). DACRs were calculated using FindAllMarkers or FindMarkers for pairwise comparisons using ‘LR’ test, min.pct of 0.05, log2 fold change threshold of 0.125, adjusted p value less than 0.05, and include the number of counts as a latent variable. ACR set enrichment was performed using AddModuleScore. Phylogenetic trees were constructed with the BuildClusterTree function. Gene activity was calculated using the GeneActivity function followed by Normalize Data with the LogNormalize method. Differentiation gene activity was calculated using FindAllMarkers or FindMarkers for pairwise comparisons using ‘LR’ test, min.pct of 0.05, log2 fold change threshold of 0.125, adjusted p value less than 0.05, and include the number of counts as a latent variable. TF motif enrichment was calculated using the functions getMatrixSet with JASPAR2020 (species 9606), CreateMotifMatrix, CreateMotifObject, and RunChromVar with BSgenome.Mmusculus.UCSC.mm10. Differentiational TF motif enrichment was calculated using FindAllMarkers or FindMarkers for pairwise comparisons using ‘LR’ test, min.pct of 0.05, log2 fold change threshold of 2, adjusted p value less than 0.05, and include the number of counts as a latent variable. Pseudotime was calculated using the Signac wrapper functions for Monocle, cluster_cells, learn_graph, order_cells, and pseudotime. The root cell was determined as max enrichment for TCF7 motif. Genome coverage tracks were generated using Signac visualization functions: CovPlot, PeakPlot, TilePlot, and AnnotationPlot.

